# ComBatFamQC: Streamlining Interactive Batch-Effect Diagnostics and Harmonization for Neuroimaging Data in R

**DOI:** 10.64898/2026.07.29.741509

**Authors:** Zheng Ren, Elizabeth A. Horwath, Siyan Wen, Randa Melhem, Jessica K. Anderson, W. Evan Johnson, Russell T. Shinohara, Andrew A. Chen, Haochang Shou

**Affiliations:** Department of Biostatistics, Bloomberg School of Public Health, Johns Hopkins University, Baltimore, MD 21205, United States; Penn Statistics in Imaging and Visualization Endeavor (PennSIVE), Department of Biostatistics, Epidemiology, and Informatics, Perelman School of Medicine, University of Pennsylvania, Philadelphia, PA 19104, United States; Center for AI and Data Science for Integrated Diagnostics (AI2D), Perelman School of Medicine, University of Pennsylvania, Philadelphia, PA 19104, United States; Department of Public Health Sciences, Medical University of South Carolina, Charleston, SC 29425, United States; 5 Division of Infectious Disease, Center for Data Science, Rutgers New Jersey Medical School, Newark, NJ 07103, United States

**Keywords:** neuroimaging harmonization, batch effect, ComBat, Shiny, diagnostics, R

## Abstract

As multisite and multi-study data aggregation becomes increasingly common for improving statistical power and sample diversity, robust harmonization methods are needed to address biases introduced by batch variation, particularly in neuroimaging research. Although a variety of harmonization approaches are available, the lack of systematic guidance for diagnosing batch effects and selecting appropriate methods remains a major challenge. To address this gap, we introduce `ComBatFamQC`, a comprehensive `R` package designed to streamline batch-effect diagnosis, harmonization, and post-harmonization analysis. `ComBatFamQC` integrates a user-friendly Shiny app for interactive batch-effect diagnostics, state-of-the-art harmonization methods from the ComBat family, including ComBat, longitudinal ComBat, ComBat-GAM, and CovBat, and tools for downstream analysis after harmonization. The package provides qualitative visualizations, statistical tests for batch-effect assessment, and a consistent interface that supports both in-sample and out-of-sample harmonization through the Shiny app, the `R` console, or the command line. In addition, it includes functions for post-harmonization analyses to facilitate downstream modeling. Its modular design also supports the systematic incorporation of future harmonization methods and expanded downstream analysis capabilities.

## 1 Introduction

Aggregating data across multiple sites and studies has become an increasingly feasible approach for enhancing statistical power and sample diversity to ensure generalizable findings. However, there remain concerns about potential bias and reduced reliability introduced by batch variations. These variations might stem from heterogeneity in study designs, data acquisition and processing pipelines across study sites. Various harmonization techniques have been applied in recent research, including ComBat and its extensions (Hu et al. 2023; Bayer et al. 2022), a commonly used statistical harmonization model to remove unwanted variation associated with sites while preserving biological associations. However, despite the existence of various harmonization techniques and software, there is a lack of clear guidance on how to diagnose batch effects and choose appropriate harmonization methods. The implementation of harmonization models remains complex and is subject to individual cases. It is often unclear as to: 1) whether a target dataset needs substantial batch correction, 2) which harmonization method to choose, and 3) whether there are any remaining significant batch effects after harmonization.

To address these questions, it is crucial to thoroughly evaluate potential batch effects for the target dataset of interest, both before and after harmonization. This process requires a series of analyses including qualitative visualization and statistical testing. We developed the `ComBatFamQC` package for the `R` environment (R Core Team 2022) to streamline such processes interactively, including batch effect diagnostics, harmonization, and post-harmonization downstream analysis. We aim to facilitate best practices to greatly simplify the data harmonization process and improve transparency and reproducibility. The package was initially developed on GitHub (https://github.com/Zheng206/ComBatFamQC) and is currently available from the Comprehensive `R` Archive Network at https://CRAN.R-project.org/package=ComBatFamQC.

Users can employ this package primarily for batch effect diagnostics and apply their own chosen harmonization techniques outside of the package. However, `ComBatFamQC` also includes four built-in ComBat Family harmonization techniques to accommodate various study designs and modeling needs. These four harmonization models are 1) ComBat, 2) longitudinal ComBat for longitudinal studies, 3) ComBat-GAM for non-linear covariate effects, and 4) CovBat for covariance batch correction with multivariate features.

The key features and overall workflow of `ComBatFamQC` are outlined below (Figure 1; Figure 2). First, it offers interactive batch effect diagnostics through a Shiny app for exploring the target dataset, and understanding the magnitude of batch variations. Additionally, it provides four commonly used harmonization techniques for in-sample and out-of-sample harmonization. After generating a harmonized dataset, users can perform various post-harmonization data preparation and downstream analyses. Currently, `ComBatFamQC` enables the creation of interactive lifespan age trend plots for individual brain features, incorporating estimated age-adjusted centiles calculated using the generalized additive models for location, scale, and shape (GAMLSS) (Rigby and Stasinopoulos 2005), with adjustments for sex and intracranial volume (ICV). These plots provide a detailed visual representation of how brain features vary across the lifespan, accounting for key covariates, and allow users to explore age-related trends and deviations interactively. `ComBatFamQC` also includes functions to easily generate covariate-corrected residuals that could be used as inputs for downstream machine learning models, facilitating robust and interpretable analyses. This paper primarily focuses on the first two aspects: *interactive batch effect diagnostics* and *harmonization*. While we continue to develop and update the post-harmonization downstream analysis functionalities, a brief example is included to illustrate the use of two post-harmonization analysis tools.

**Figure 1:**
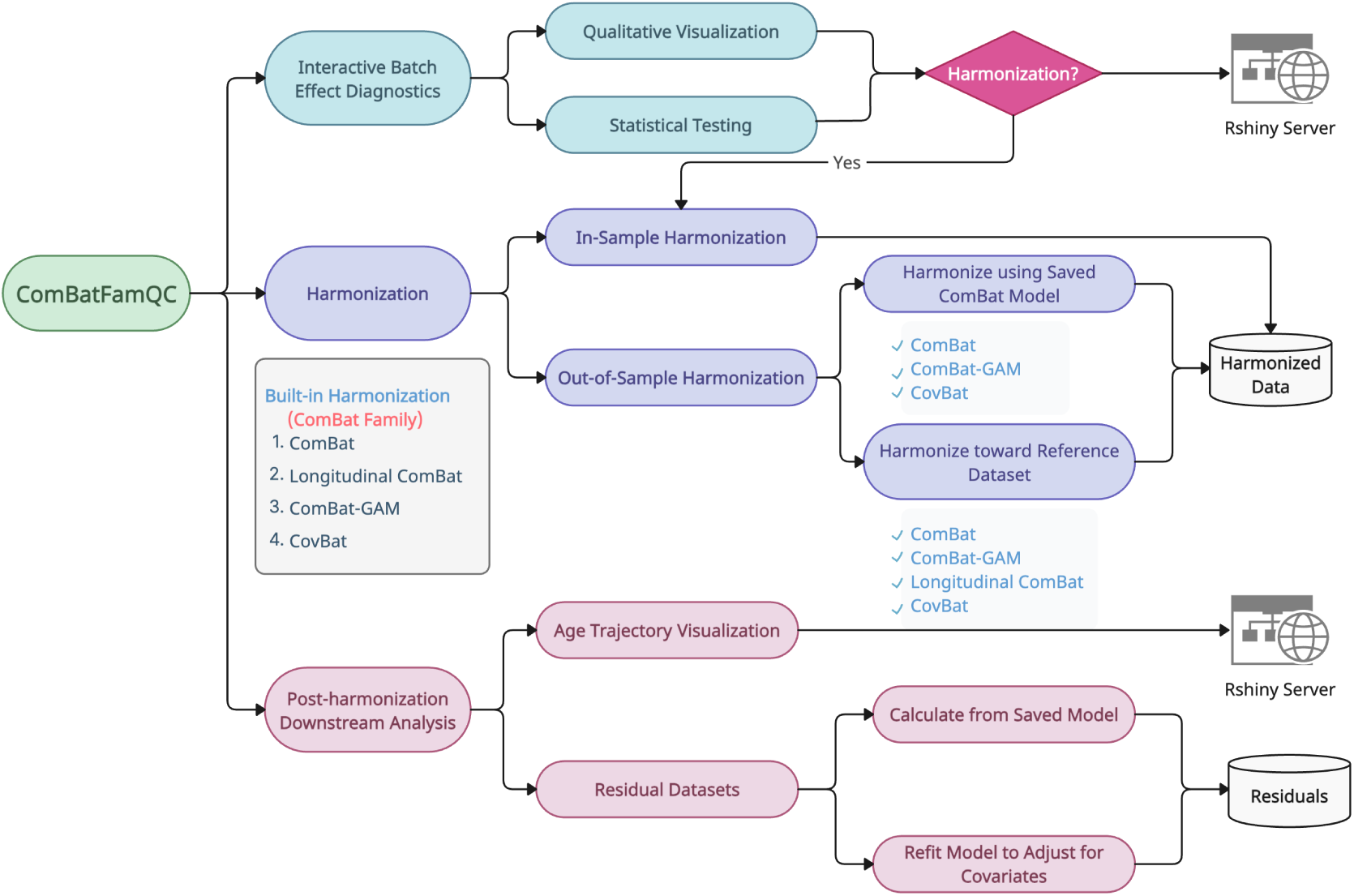
Overview of the `ComBatFamQC` framework. This schematic summarizes the main components of `ComBatFamQC`, including interactive batch effect diagnostics, harmonization, and post-harmonization downstream analysis. The package provides four built-in harmonization methods from the ComBat family (ComBat, Longitudinal ComBat, ComBat-GAM, and CovBat) and supports both in-sample and out-of-sample harmonization. It also includes two Shiny applications: one for interactive batch effect diagnostics and harmonization, and another for age trajectory visualization and residual-based downstream analyses.

**Figure 2:**
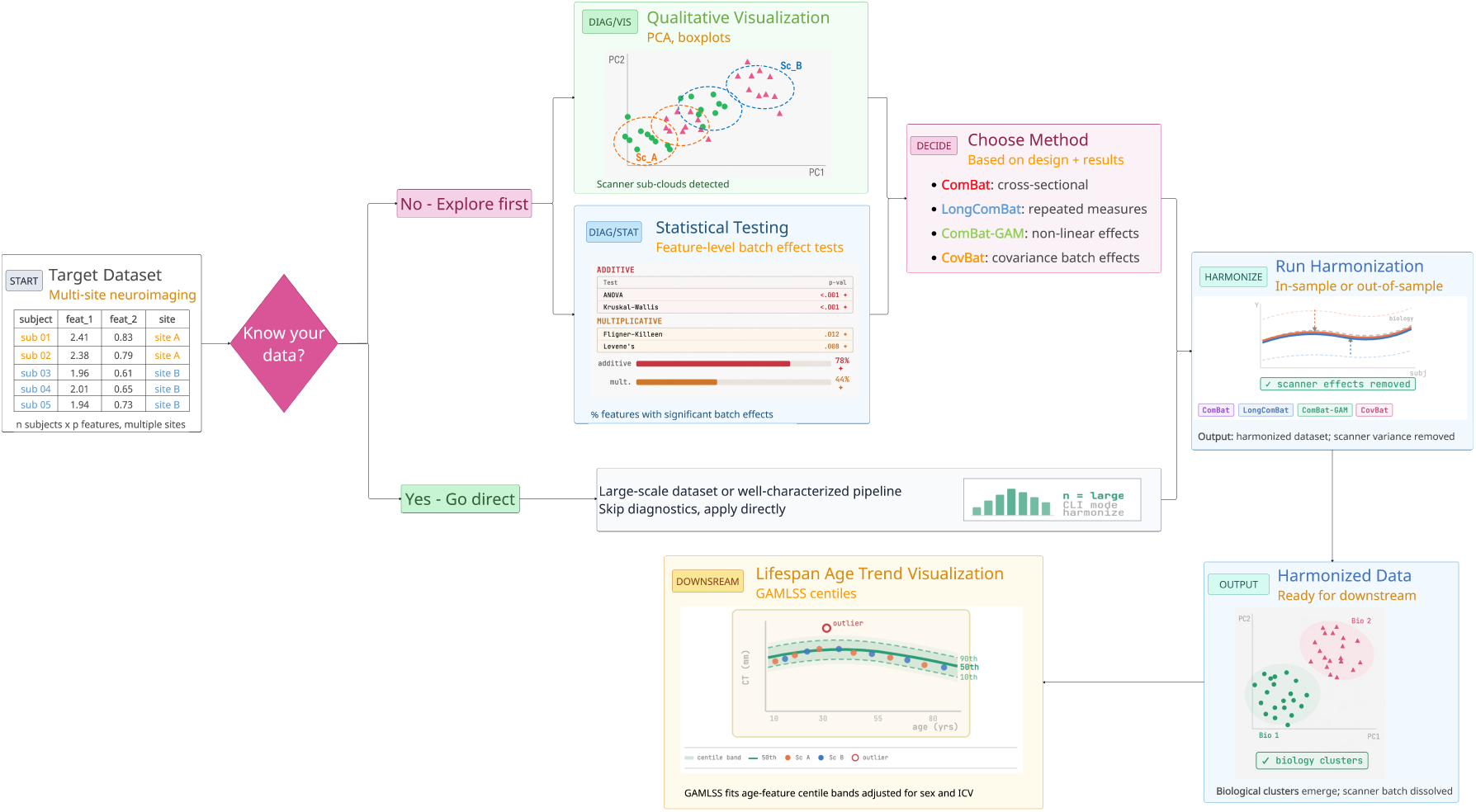
Illustrative user workflow in `ComBatFamQC`. This example shows how a user may move through the package starting from a multi-site neuroimaging dataset. Users can either follow an exploratory path, in which batch effects are examined through interactive visualizations and feature-level statistical tests to inform method selection, or proceed directly to harmonization when the dataset and acquisition protocol are already well characterized. After harmonization, the output dataset can be used for downstream analyses, such as interactive lifespan age-trend visualization based on GAMLSS centile curves adjusted for sex and intracranial volume (ICV).

The rest of the paper is organized as follows. Section 2 provides an overview of both the ComBat Family methods and batch effect diagnostics methods to help users better understand the package. Section 3 introduces the key features of the package, and Section 4 presents results from an example application to the ADNI dataset. We close in section 5 with concluding remarks.

## 2 Methods

### 2.1 ComBat Family Methods Overview

In this section, we first provide an overview of the methods behind the four ComBat Family harmonization techniques integrated into the `ComBatFamQC` package.

#### 2.1.1 Original ComBat

The original ComBat method (W. E. Johnson, Li, and Rabinovic 2006; Fortin, Parker, et al. 2017; Fortin, Cullen, et al. 2018) employs an empirical Bayes (EB) linear model framework and explicitly models data variations attributed to covariates, and additive and multiplicative batch effects. Briefly, ComBat assumes that each observed imaging feature *y_ijv_*, such as cortical thickness or volume measures, follows

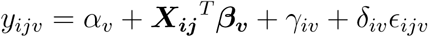

where *i* is the batch index that could represent study, site, or scanner as defined by user, *j* represents subject within batch, *v* is the feature index. The statistical associations between *y_ijv_* and covariate ***X_ij_*** are modeled using a linear regression framework where ***β_v_*** is the feature-specific slope and *α_v_* is the intercept. In addition, *γ_iv_* accounts for the batch-specific shift in feature-mean (additive batch effect) and *δ_iv_* accounts for variance differences due to batch (multiplicative batch effect). *ɛ_ijv_* is the error term and is assumed to independently follow *N* 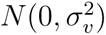. In the first step, ComBat obtains the least-squares estimates *α̂_v_* and ***β̂v*** for each feature. ComBat then assumes that the site parameters follow a common prior distribution within batch across features such that *γ_iv_* follows independent normal distributions and *δ_iv_* follows independent inverse gamma distributions. The empirical Bayes method is employed to obtain the posterior estimate of the hyperparameters 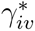 and 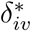, which are then removed from the residuals. The ComBat-harmonized data could be expressed as:

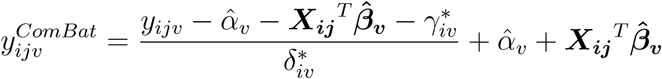

#### 2.1.2 ComBat-GAM

ComBat-GAM (Pomponio et al. 2020) extends the original ComBat model by allowing for flexible non-linear covariate effects via the generalized additive model (GAM). ComBat-GAM has been successfully applied to harmonizing multi-site datasets with large age range over lifespan. ComBat-GAM model could be expressed as:

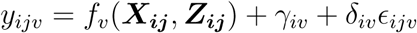

where

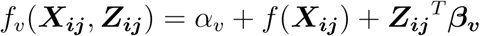

is the nonlinear covariate model. *f* (***X_ij_***) is a nonlinear model for ***X_ij_***, based on splines, while ***Z_ij_*** represents the linear covariates, and ***β_v_*** is the linear slope. *f_v_*(***X_ij_***, ***Z_ij_***) denotes the variation of feature *v* value captured by the biologically-relevant covariates. The batch parameters *γ_iv_* and *δ_iv_* could be estimated similarly to ComBat after removing the covariate effects. The ComBat-GAM-harmonized data can be expressed as:

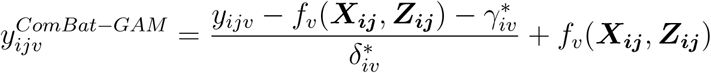

where 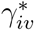 and 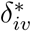 are the empirical Bayes estimates under the same assumptions as the original ComBat method.

#### 2.1.3 Longitudinal ComBat

For imaging data collected from longitudinal studies, it is necessary to account for within-subject correlations among the features. Longitudinal ComBat (LongComBat) (Beer et al. 2020) was designed to effectively harmonize longitudinal observations and repeated measures from traveling subjects studies. By explicitly including the subject-specific random intercept and time variable and combining linear mixed effects regression with ComBat model, the longitudinal ComBat can be expressed as:

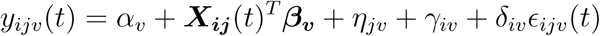

Compared with the cross-sectional ComBat model, the covariate design matrix ***X_ij_***(*t*) includes a time variable and potential interaction terms with time. *η_jv_* is a subject-specific random intercept that accounts for within-subject correlation. *γ_iv_* is the additive batch effect and *δ_iv_* is the multiplicative batch effect. The LongComBat-harmonized data can be expressed as:

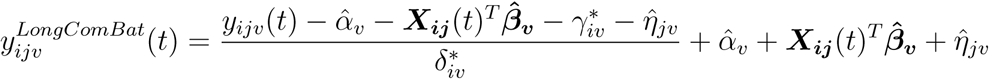

where *α̂_v_*, ***β̂v*** and *η̂_jv_* are the fixed effect parameters estimated by the best linear unbiased estimator (BLUE), and 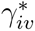 and 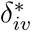 are the empirical Bayes estimates under the same assumptions as above.

#### 2.1.4 CovBat

Lastly, Correcting Covariance Batch Effects (CovBat) (Chen et al. 2021) was developed to address situations where batch effects are present not only in individual features but also in the covariance structure between features. Covariance captures the joint relationships between features, which are crucial for machine learning (ML)-based approaches in neuroimaging. Variations in covariance across sites can distort these relationships, resulting in biased comparisons, the masking of biologically meaningful associations, and the introduction of spurious findings. Thus, addressing batch effects in covariance is essential for ensuring robust and reproducible multi-site ML-based analyses.

CovBat can be viewed as a two-stage harmonization procedure. In the first stage, standard ComBat is used to remove batch-related mean and variance shifts in the marginal distributions of the target data, yielding ComBat-adjusted residuals:

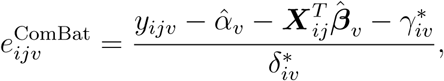

where 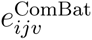 denotes the ComBat-adjusted residual, which is assumed to have mean zero and batch-specific covariance matrix **Σ***_i_*. Here, *α̂_v_*, ***β̂****v*, 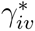, and 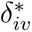 are estimates from the original ComBat model.

Let

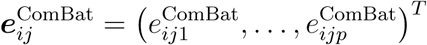

denote the vector of ComBat-adjusted residuals across all *p* features for subject *j* in batch *i*. In the second stage, CovBat harmonizes the ComBat-adjusted residuals in principal component space by shifting batch-specific covariance structures toward the population-average covariance structure:

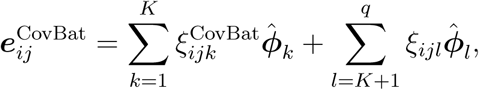

where 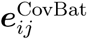 is the residual vector after covariance harmonization, *ξ_ijl_* is the original principal component score, 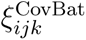 is the harmonized principal component score, *K* is the number of principal component scores to be harmonized, and *q* is the total number of principal components. The vectors ***ϕ̂****k* and ***ϕ̂****l* denote the estimated eigenvectors.

Finally, CovBat reconstructs the harmonized data by adding back the preserved intercept and covariate effects:

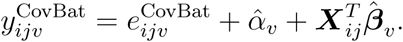

This reconstruction step preserves covariate-associated biological variation while producing CovBat-adjusted observations.

### 2.2 Batch Effect Diagnostics

We then delve into a range of visualization and statistical testing methods integrated into the package to evaluate possible batch effects in the target dataset. These evaluation methods can be broadly categorized into two main types (Table 1): 1) **Feature-level Evaluation**, and 2) **Global Evaluation**. The Feature-level Evaluation is designed to assess feature-specific batch variations in mean or variance, while the Global Evaluation aims to evaluate whether potential batch effects are present in the entire dataset, encompassing all features.

**Table 1:**
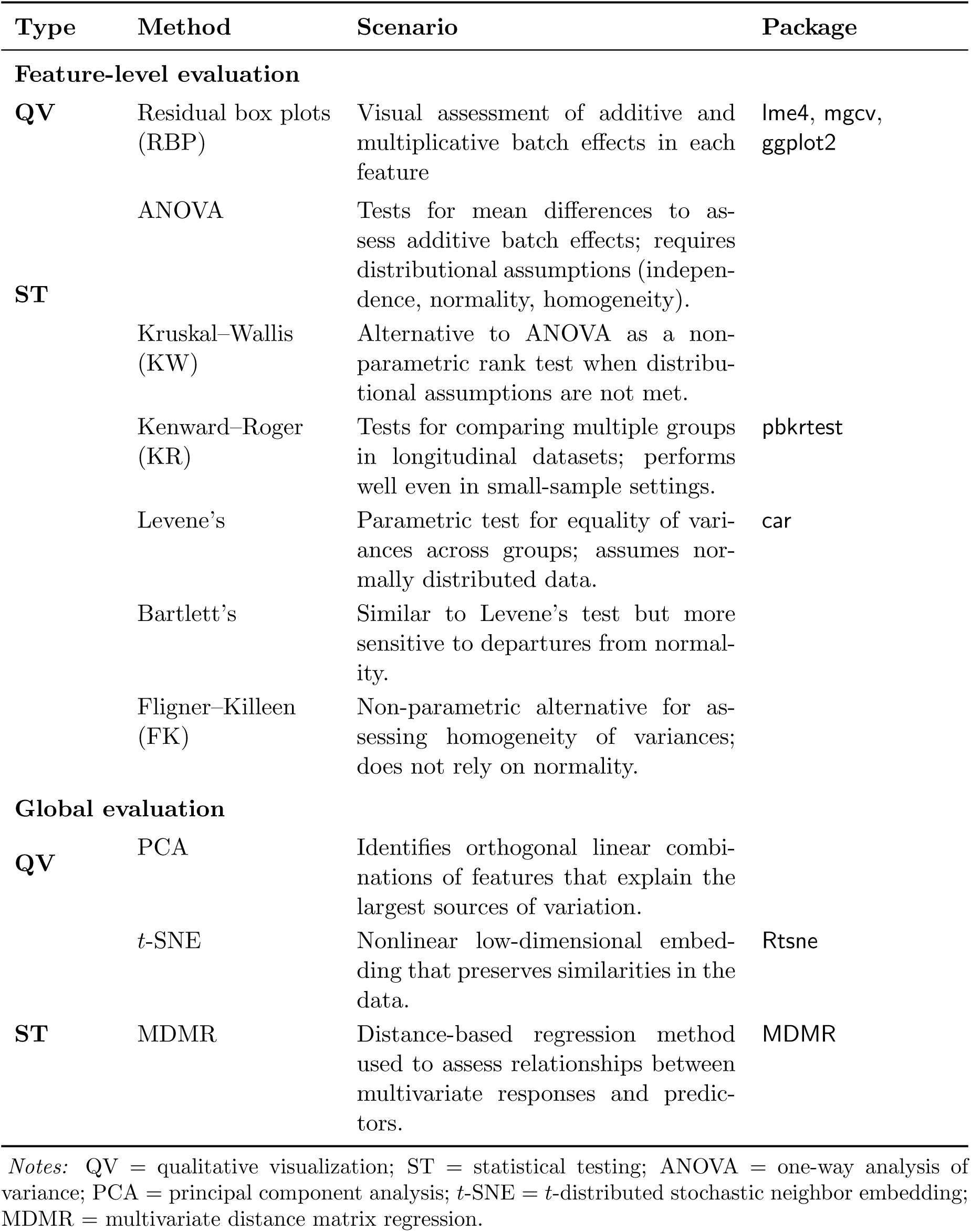
Batch effect diagnostics method summary.

#### 2.2.1 Feature-level Evaluation

##### a. Qualitative Visualization

Our package includes two types of **residual box plots (RBP)** for visual assessment of additive and multiplicative batch effects in each feature: 1) **additive-residual box plots** and 2) **multiplicative-residual box plots**. A noticeable deviation of the mean from zero in the additive-residual box plot indicates the presence of an additive batch effect, whereas a substantial variation in the variance of the multiplicative-residual box plot implies a potential multiplicative batch effect.

The residuals used to plot both types of box plots were derived by estimating and removing the variations associated with model covariates through appropriate regression models. Take cross-sectional data as an example, we first fit the following model where each feature serves as the dependent variable, with preserved covariates and the batch variable as predictors:

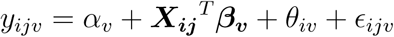

where *i* is the batch index, *j* is the participant index, *v* is the feature index, *y_ijv_*is the observed value of feature v, *α_v_* is the intercept, ***X_ij_*** is a vector of covariates to be adjusted, and ***β_v_*** is the vector of regression coefficients corresponding to ***X_ij_***, *θ_iv_* corresponds to the additive mean shift of batch level *i* for feature *v* and *ɛ_ijv_*is the error term.

Residuals to examine additive batch effect are subsequently computed as:

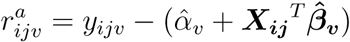

Residuals for evaluating multiplicative batch effect are computed by removing both the variability associated with model covariates and the estimated additive batch effects as:

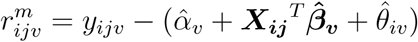

where *α̂_v_*, ***β̂v*** and *θ̂iv* are least-squares estimates. Note that the choice of the regression models can be tailored to suit specific study designs. Within this package, users could choose to fit a 1) linear model (lm), 2) linear mixed-effect model from the `R` package `lme4` (Bates et al. 2015) (lmer), and 3) generalized additive model from the `R` package `mgcv` (Wood 2011) (gam). Box plots are generated using the `R` package `ggplot2` (Wickham 2016).

##### b. Statistical Testing Methodology

In addition to qualitative visualizations, we implemented several statistical tests to quantitatively evaluate additive and multiplicative batch effects separately.

###### Additive Batch Effects

Three statistical tests for mean differences are included to assess additive batch effects: one-way analysis of variance (ANOVA), Kruskal-Wallis (KW), and Kenward-Roger (KR) tests. A significant test statistic suggests the potential presence of an additive batch effect within the specific feature.

**one-way ANOVA** calculates p-values based on several distributional assumptions of the data including the independence of observations, the normality of residuals, and the homogeneity of variance.

When the residuals are not normally distributed or the equal variance assumption is violated, the **Kruskal-Wallis (KW)** test (Kruskal and Wallis 1952) is a more appropriate alternative to ANOVA as a non-parametric rank test for comparing differences between two or more groups.

Lastly, the **Kenward-Roger (KR)** (Kenward and Roger 1997) test is included for comparing multiple batches in longitudinal imaging features. This method has been demonstrated to perform well, particularly in small sample settings, as shown in the original study. The `ComBatFamQC` package integrates the KR test through `R` package `pbkrtest` (Halekoh and Højsgaard 2014).

###### Multiplicative Batch Effects

We used statistical tests for variance differences to detect multiplicative batch effects, which include Levene’s, Bartlett’s and Fligner-Killeen (FK) tests. A significant statistic suggests the potential existence of a multiplicative batch effect within the specific feature.

**Levene’s** test (Levene et al. 1960) and **Bartlett’s** test (Bartlett 1937) are both parametric tests for assessing the equality of variances across multiple groups. They both assume that the data are normally distributed and the Bartlett’s test is more sensitive to departures from normality. The `ComBatFamQC` package integrates the Levene’s test through the `R` package `car` (Fox and Weisberg 2019).

We also included the **Fligner-Killeen (FK)** (Conover, M. E. Johnson, and M. M. Johnson 1981) test as a non-parametric alternative to Levene’s and Bartlett’s tests for assessing the homogeneity of variances. This test is particularly robust when parametric assumptions, such as normality, are violated, making it a valuable choice for datasets with non-normal distributions.

#### 2.2.2 Global Evaluation

##### a. Qualitative Visualization

**Principal component analysis (PCA)** (Hotelling 1933) identifies orthogonal linear combinations of the original features that represent the most significant variations in the data. We applied PCA on residuals with additive batch effects, as described in Section 2.2.1, and included an interactive scatter plot featuring any selected pair of principal components. The scatter plot is color-coded based on the batch variable, and the presence of a discernible color pattern that facilitates clustering indicates the persistence of global batch effects.

**T-distributed stochastic neighbor embedding (***t***-SNE)** is a nonlinear lower dimensional embedding technique used for data visualization (van der Maaten and Hinton 2008). It constructs a probability distribution that represents pairwise similarities between data points and aims to find a lower-dimensional representation of the data that best preserves these similarities. We applied *t*-SNE method on residuals with additive batch effects and included a scatter plot colored by the batch variable. Similarly to PCA plots, the presence of any discernible color pattern that helps with clustering suggests the existence of global batch effects. The `ComBatFamQC` package integrates the *t*-SNE method through `R` package `Rtsne` (Jack et al. 2008).

##### b. Statistical Testing Methodology

Beyond qualitative visualizations, **multivariate distance matrix regression (MDMR)** (McArdle and Anderson 2001) is utilized to evaluate the overall presence of batch effects among datasets. MDMR is a distance-based regression method used to assess the relationships between multivariate response variables and a set of predictor variables. It considers the dissimilarity matrices of the response variables and aims to determine how well the predictor variables explain the variation in the responses, while accounting for the similarities and differences among data points. A significant coefficient for the batch effect in MDMR suggests the presence of global batch effects. The `ComBatFamQC` package integrates the MDMR method through the `R` package `MDMR` (McArtor, Lubke, and Bergeman 2016). Note that MDMR is based on permutation and can be computationally intensive for large datasets. `ComBatFamQC` provides the option to disable the MDMR test by setting the parameter *mdmr* to FALSE.

## 3 Features of ComBatFamQC

As mentioned above, `ComBatFamQC` streamlines batch-effect diagnostics, harmonization, and post-harmonization downstream analyses. In this section, we summarize the key functionality provided by the package. Datasets used for batch-effect diagnostics and harmonization should include (i) a batch variable, (ii) feature column(s) to be harmonized, (iii) optional covariate column(s) to be adjusted in the mean model, and (iv) random-effect column(s) for longitudinal data.

### 3.1 Interactive Batch Effect Diagnostics

`ComBatFamQC` supports interactive batch-effect diagnostics through a Shiny-based workflow that integrates qualitative visualization and statistical testing. The diagnostics are conducted at three complementary levels: (i) exploratory summaries of batch composition and covariate distributions to assess potential confounding, (ii) feature-level diagnostics targeting additive and multiplicative batch effects, and (iii) global diagnostics that evaluate multivariate batch effects across the full feature set. Example diagnostic outputs are presented in the Results section, while interface screenshots are provided in the Supplementary Material.

Preparation for this workflow is handled by the `visual prep` function, which accepts a customizable set of parameters and computes the diagnostic quantities and statistical test results described in Section 2.2. In addition to generating diagnostic outputs, `visual prep` returns summary statistics and user-specified metadata (e.g., batch and covariate names), enabling users to verify that variables have been correctly specified. A detailed description of the function parameters is provided in Table 2.

**Table 2:** Batch-effect diagnostics–related parameters for visual prep.

| Parameter | Short description |
| --- | --- |
| `df` | Target dataset to be evaluated. |
| `type` | Regression model used to describe covariate-feature associations: “lm”, “lmer”, or “gam” (“lm” by default). |
| `features` | Column names of the features to be harmonized. |
| `batch` | Column name of the batch variable. |
| `covariates` | Covariates to be preserved (NULL by default). |
| `interaction` | Interaction terms to include (e.g., “cov1,cov2”; NULL by default). |
| `smooth_int_type` | Interaction type used in the “gam” model: “linear”, “categorical-continuous”, “factor-smooth”, “tensor”, or “smooth-smooth” (“linear” by default). |
| `smooth` | Covariates modeled nonlinearly (NULL by default). |
| `random` | Random-effect variable for mixed models (NULL by default). |
| `cores` | Number of CPU cores for parallel computation (`detectCores()` by default; for **Windows**, set `cores = 1`). |
| `mdmr` | Whether to disable the MDMR global diagnostic to reduce computation time (TRUE by default). |

The output of `visual_prep` is then passed to `comfam_shiny`, which launches the interactive diagnostics module. The argument *after* determines whether diagnostics are applied to the original dataset before harmonization (*after = FALSE*) or to the harmonized data for post hoc evaluation (*after = TRUE*).

The interactive diagnostics module is organized into five distinct tabs, each addressing an important aspect of batch-effect diagnostics for the target data. The first two tabs focus on data exploration, while the remaining three address different diagnostic components.

The **Data Overview** section provides users with an overview of the data they uploaded (Figure S1), highlighting the batch column, covariates columns, and features columns in blue, pink, and light yellow, respectively. This allows users to determine if they have specified the columns correctly. Additionally, this section provides simple exploratory data analysis through graphical tools, allowing users to explore specific features and investigate how their distributions vary by batch levels and their associations with potential covariates. These tools include density plots of features stratified by batch, box plots of categorical covariates across batch levels, and scatter plots with fitted trend lines to examine bivariate relationships between two continuous variables (Figure S2).

The **Summary** tab provides both tabular summary statistics and graphical displays, including bar plots and box plots, to illustrate the sample size distribution of the batch variable and the distributions of covariates (Figure S3). In the batch sample size summary table, a `remove` column indicates whether specific batch levels have been excluded from the dataset based on their sample size (Figure S4). In practice, batch levels with fewer than three observations are typically removed to avoid failures in statistical modeling.

The **Residual Plot** section provides qualitative visualizations to help users assess potential batch effects for individual features. This tab displays box plots of residuals for a selected feature after removing variation associated with covariates, allowing users to evaluate possible feature-level additive and multiplicative batch effects (Figure S5). The residual distributions are stratified by batch, with batch levels displayed on the x-axis. In the additive residual plots, deviations of the residual means from zero or systematic differences across batch levels suggest additive batch effects. In the multiplicative residual plots, substantial differences in interquartile ranges across batch levels indicate multiplicative batch effects. Users can explore different features using the drop-down menu in the left panel.

The **Diagnosis of Global Batch Effect** section provides qualitative visualizations of batch-related clustering through PCA and *t*-SNE, along with a quantitative assessment based on the MDMR model. In the PCA and *t*-SNE plots, points are colored by batch, and clear separation by color suggests the presence of global batch effects across features (Figure S6). Users can select which principal components to display in the PCA plot, and the corresponding proportion of variance explained is reported in the table below. The MDMR table summarizes the statistical test results for global batch effects. A highlighted row with a p-value less than 0.05 indicates a significant batch effect across the feature set.

The **Diagnosis of Feature-level Batch Effect** section presents detailed statistical test results for batch effects on individual features. As noted earlier, diagnostics for additive batch effects include the results of the ANOVA, Kruskal–Wallis, and Kenward– Roger tests, with the latter available only for linear mixed models. Diagnostics for multiplicative batch effects include the results of the Fligner–Killeen, Levene’s, and Bartlett’s tests. Significant results are highlighted in the tables, and below each table, a summary is provided showing the percentage of features with significant batch effects out of the full feature set (Figure S7).

Together, these five tabs provide users with a thorough understanding of the target datasets and help them evaluate the extent of any remaining need for harmonization. After completing the diagnostic review, users may export all results as either a combined Excel file or a Quarto report, depending on their preference and whether Quarto is installed, thereby completing the quality control process.

### 3.2 Harmonization

In addition to batch-effect diagnostics, `ComBatFamQC` integrates four methods from the ComBat family, enabling harmonization through both an interactive workflow in the Shiny app and programmable interfaces, including R and the command line. The interactive workflow is provided within the same Shiny app as the batch diagnostic module through the “Interactive Harmonization” tab, which is available following the diagnostic session. This interface enables users to perform harmonization using built-in ComBat family methods without writing custom code (Figure S8). Within this tab, users can specify harmonization parameters, select an appropriate ComBat model, run an empirical Bayes (EB) assumption test, and define a file path for saving both the harmonized dataset and the fitted ComBat model. Equivalent functionality is also available through the R console or through a controlled command-line pipeline for users with more advanced programming experience. Overall, the interactive workflow is designed to support fast, guided harmonization through a graphical user interface informed by diagnostic results, whereas the command-line interface is better suited to scalable analyses and programmatic workflows.

The harmonization procedure is implemented through `combat_harm`, which returns both the harmonized dataset and the fitted harmonization model. The function supports two common use cases: (i) *in-sample* harmonization, in which a dataset is harmonized directly after acquisition using all available data, and (ii) *out-of-sample* harmonization for ongoing multisite studies, in which model estimates learned from a training dataset are applied to newly acquired data (Xin et al. 2026). Out-of-sample harmonization may be performed either by applying an existing fitted model to new data or by aligning the new data to an existing reference dataset.

When an existing model is applied, `combat_harm` compares the batch variables in the new dataset with those used to train the harmonization model. If the new data contain batch levels observed during model training, the corresponding batch-effect estimates are applied directly. For previously unseen batches, batch-effect parameters are estimated using the empirical Bayes framework, under the assumption that the same mean model and prior distributions remain appropriate.

In contrast, reference-based harmonization assumes that batch effects have already been fully removed from the reference dataset, which is then treated as originating from a single reference site. The remaining batches are subsequently aligned to this reference level. Currently, the application of existing fitted models is supported for linear models and generalized additive models (GAMs), but not for linear mixed-effects models, due to the difficulty of predicting unseen random effects. Consequently, reference-based harmonization offers a practical alternative for longitudinal data settings.

Because all ComBat-based methods rely on empirical Bayes shrinkage, assessing the validity of the EB assumption is an important part of the harmonization process. To facilitate this, `ComBatFamQC` provides an EB assumption-checking feature that allows users to compare the prior distribution with the empirical distribution through the interactive Shiny app. The same feature can be enabled by setting `eb_check = TRUE` when calling `combat_harm`. As part of this assessment, line plots of the estimated location and scale parameters are generated to help users determine whether the EB assumption is reasonable. In these plots, solid lines denote the EB-based prior distributions, whereas dotted lines denote the empirical density estimates. In general, substantial overlap between the two sets of lines in both plots indicates that the EB assumption is reasonably satisfied and that EB-based harmonization can be used appropriately.

Notably, the `combat_harm` function shares several parameters with `visual_prep` that describe the data structure, including the batch variable, the features to be harmonized, covariates, and interaction terms. Detailed descriptions of the additional harmonization parameters are provided in Table 3.

**Table 3:** Harmonization-related parameters.

| Parameter | Short description |
| --- | --- |
| eb_check | Whether to run the empirical Bayes (EB) assumption check or perform harmonization (`FALSE` by default). |
| family | Harmonization model family: ComBat or CovBat (`comfam` or `covfam`; `comfam` by default). |
| type | Harmonization model type: “lm”, “lmer”, or “gam” (“lm” by default). |
| eb | Whether to use empirical Bayes estimation for mean and variance (`TRUE` by default). |
| ref.batch | Reference batch level (`NULL` by default). |
| predict | Whether to use an existing model for out-of-sample harmonization (`FALSE` by default). |
| object | Previously fitted ComBat model object (`NULL` by default). |
| reference | Reference dataset for out-of-sample harmonization (`NULL` by default). |
| out_ref_include | Whether to include the reference dataset in the harmonized output (`TRUE` by default). |

**Table 4:** Batch-effect diagnostic results before and after harmonization.

| Method | unharm | harm |
| --- | --- | --- |
| ANOVA | 46.77% | 0% |
| Kruskal–Wallis | 100% | 29.03% |
| Kenward–Roger | 48.39% | 0% |
| Levene | 93.55% | 1.61% |
| Bartlett | 98.39% | 0% |
| Fligner–Killeen | 95.16% | 9.68% |
| MDMR | < 0.001 | < 0.001 |

### 3.3 Post-Harmonization Downstream Analysis

In addition to batch-effect diagnostics and harmonization, `ComBatFamQC` provides two tools to support post-harmonization downstream analyses: (1) customized residual generation and (2) lifespan age trend visualization.

#### a. Customized Residual Generation

The package allows users to generate residuals from harmonized data to facilitate down-stream analyses that require selective removal of covariate effects. This functionality is implemented through the `residual gen` function, which supports flexible model specification via parameters controlling model type, covariates, interaction terms, random effects, and smoothing options. Residuals may be generated either by fitting a new regression model or by reusing an existing fitted model, enabling efficient and reproducible analyses.

#### b. Lifespan Age Trend Visualization

`ComBatFamQC` also includes tools for investigating lifespan age trends in brain features while adjusting for relevant covariates such as sex and intracranial volume (ICV). Age trend estimation is implemented via the `age list gen` function, which fits generalized additive models for location, scale, and shape (GAMLSS) to estimate age-dependent centiles for individual features. Detailed descriptions of the function parameters are provided in Table S1.

To facilitate interactive exploration, the `age shiny` function launches a Shiny application that visualizes observed feature values alongside estimated age trends at selected centiles. The app also provides tabular summaries of estimated values across ages and allows users to customize visualization settings, select centile levels, and export both the estimated trends and fitted models for further analysis.

## 4 Results

### 4.1 Data

To demonstrate the application and usage of the `ComBatFamQC` package, we use publicly available longitudinal cortical thickness data from the Alzheimer’s Disease Neuroimaging Initiative (ADNI) study (Jack et al. 2008). ADNI was approved by the institutional review boards of all participating institutions, and written informed consent was obtained from all participants or their authorized representatives. The present work involved secondary analysis of publicly available, de-identified data and did not involve new data collection from human participants.

The dataset contains cortical thickness measures from 62 brain regions for 663 unique participants (282 females and 381 males; mean baseline age 75 *±* 6.7 years), with images collected longitudinally across 2–6 visits. The data were processed and are available for download at https://github.com/ntustison/CrossLong. The dataset is also included in the package as sample data.

In this dataset, scanner manufacturer is considered the primary batch variable, with three categories: Siemens, Philips, and GE. As shown in Figure 3A, Siemens and GE scanners are more frequently represented than Philips. The covariates are distributed relatively evenly across batch levels, suggesting minimal confounding between covariates and the batch variable. Across all batch levels, a larger proportion of participants are diagnosed with late mild cognitive impairment (LMCI), and males are more prevalent than females.

**Figure 3:**
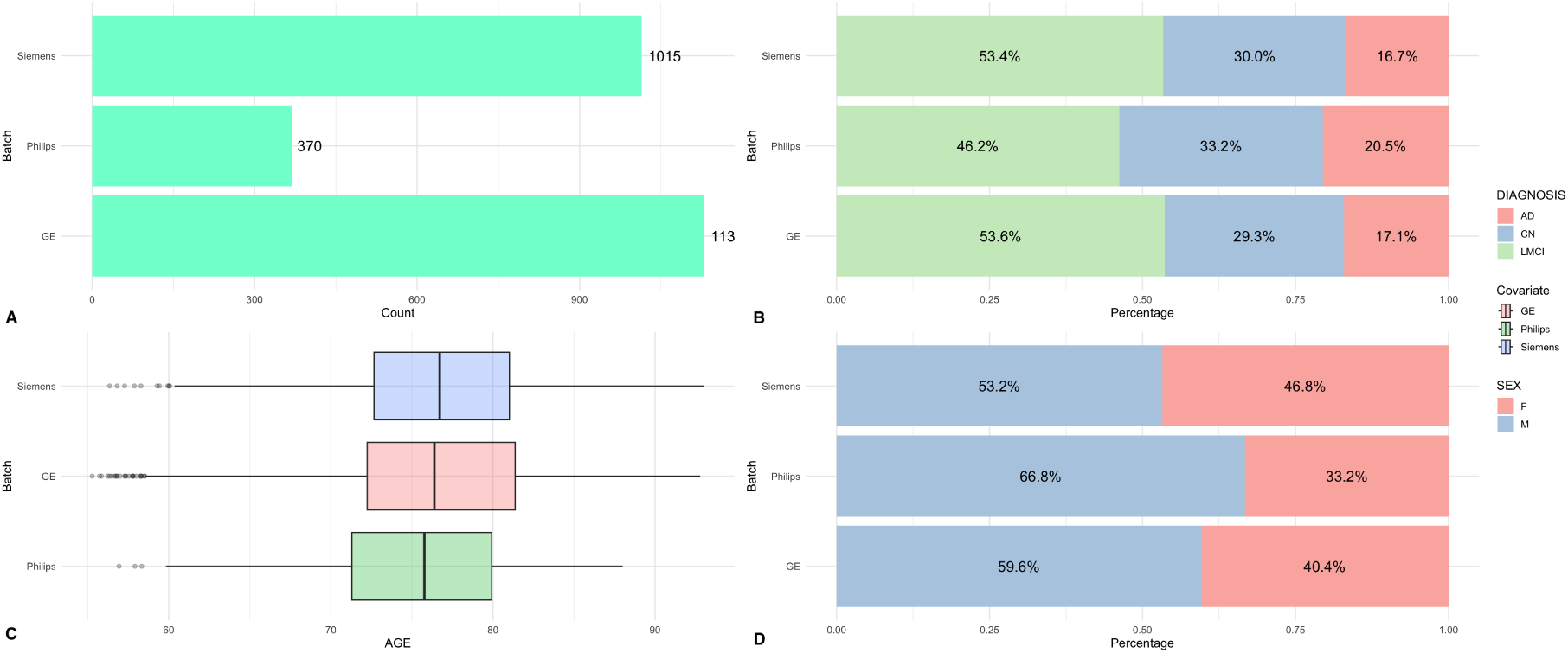
Exploratory data analysis of the ADNI cortical thickness dataset. (A) Distribution of scanner manufacturer across observations. (B–D) Distribution of key covariates across scanner manufacturers. Siemens and GE scanners are more frequently represented than Philips, while covariates appear to be relatively evenly distributed across batch levels.

### 4.2 Batch Effect Diagnostics

We first examined qualitative visualizations to assess feature-level batch effects. For the representative feature *thickness.left.lateral.orbitofrontal*, both the feature distribution plot (Figure 4A) and the age-trend plot (Figure 4D) showed a clear shift in mean across batch levels, indicating the presence of batch-related differences. The additive residual plot provided further evidence of pronounced additive batch effects, whereas the multiplicative residual plot suggested comparatively weaker evidence of multiplicative batch effects for this ROI (Figure 4B,E). In addition, both the PCA and *t*-SNE plots revealed clear global batch effects across features (Figure 4C,F).

**Figure 4:**
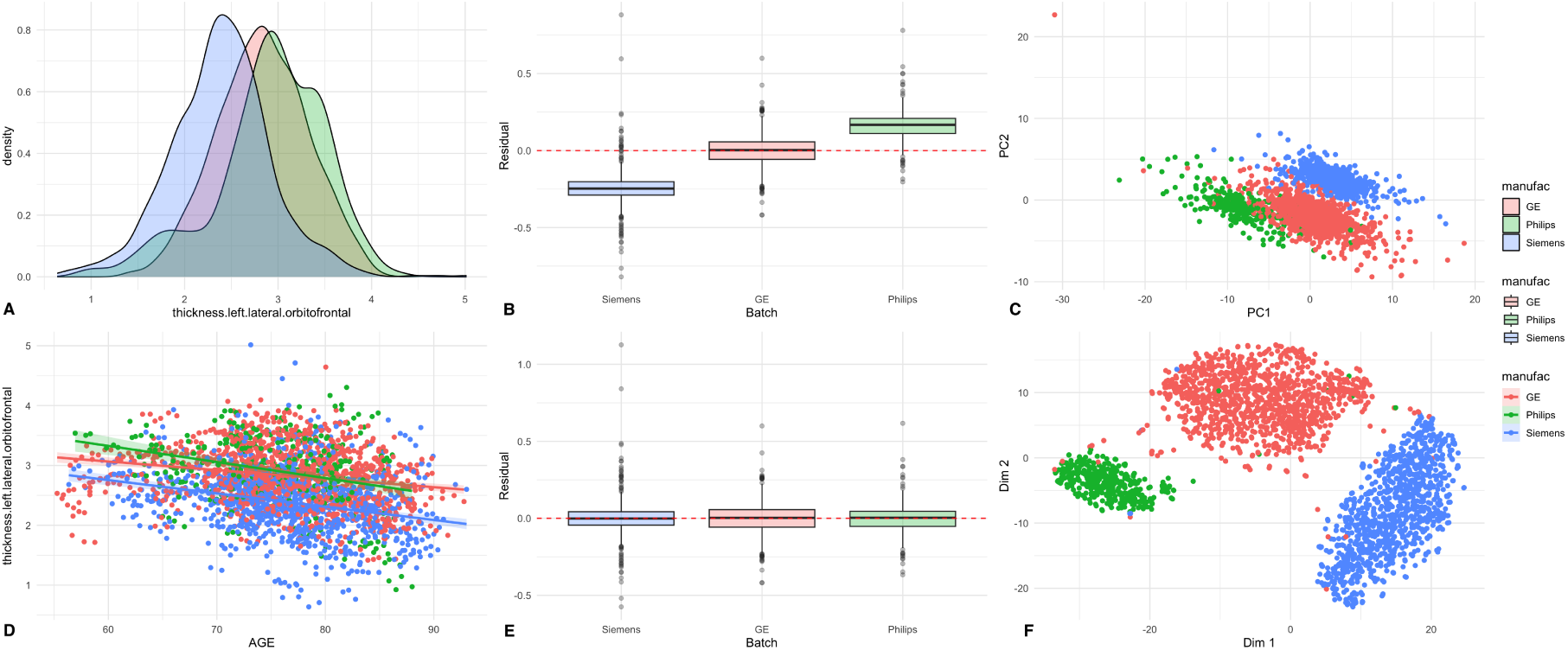
Pre-harmonization batch diagnostics. For *thickness.left.lateral.orbitofrontal*, the feature distribution (A) and age-trend plot (D) show mean shifts across batches. The additive residual plot (B) suggests stronger additive effects, whereas the multiplicative residual plot (E) suggests weaker multiplicative effects. The PCA plot (C) and t-SNE plot (F) show separation by batch at the multivariate level.

These visual findings were supported by formal statistical testing at the feature level. Specifically, 48.39% of ROIs showed significant additive batch effects according to the Kenward-Roger test, while 95.16% exhibited significant multiplicative batch effects based on the Fligner-Killeen test. The ANOVA results were consistent with those of the Kenward-Roger test, whereas the Kruskal-Wallis test appeared overly sensitive, identifying 100% of ROIs as significant. Similarly, Levene’s and Bartlett’s tests produced results comparable to those of the Fligner-Killeen test.

Beyond the feature level, the MDMR analysis yielded a p-value of less than 0.001, indicating strong global batch effects across features and providing clear justification for harmonization using longitudinal ComBat. We also performed an EB assumption check to evaluate the suitability of the empirical Bayes approach. The resulting line plots showed close agreement between the prior and empirical distributions, suggesting that the EB assumptions were reasonably well satisfied (Figure 5).

**Figure 5:**
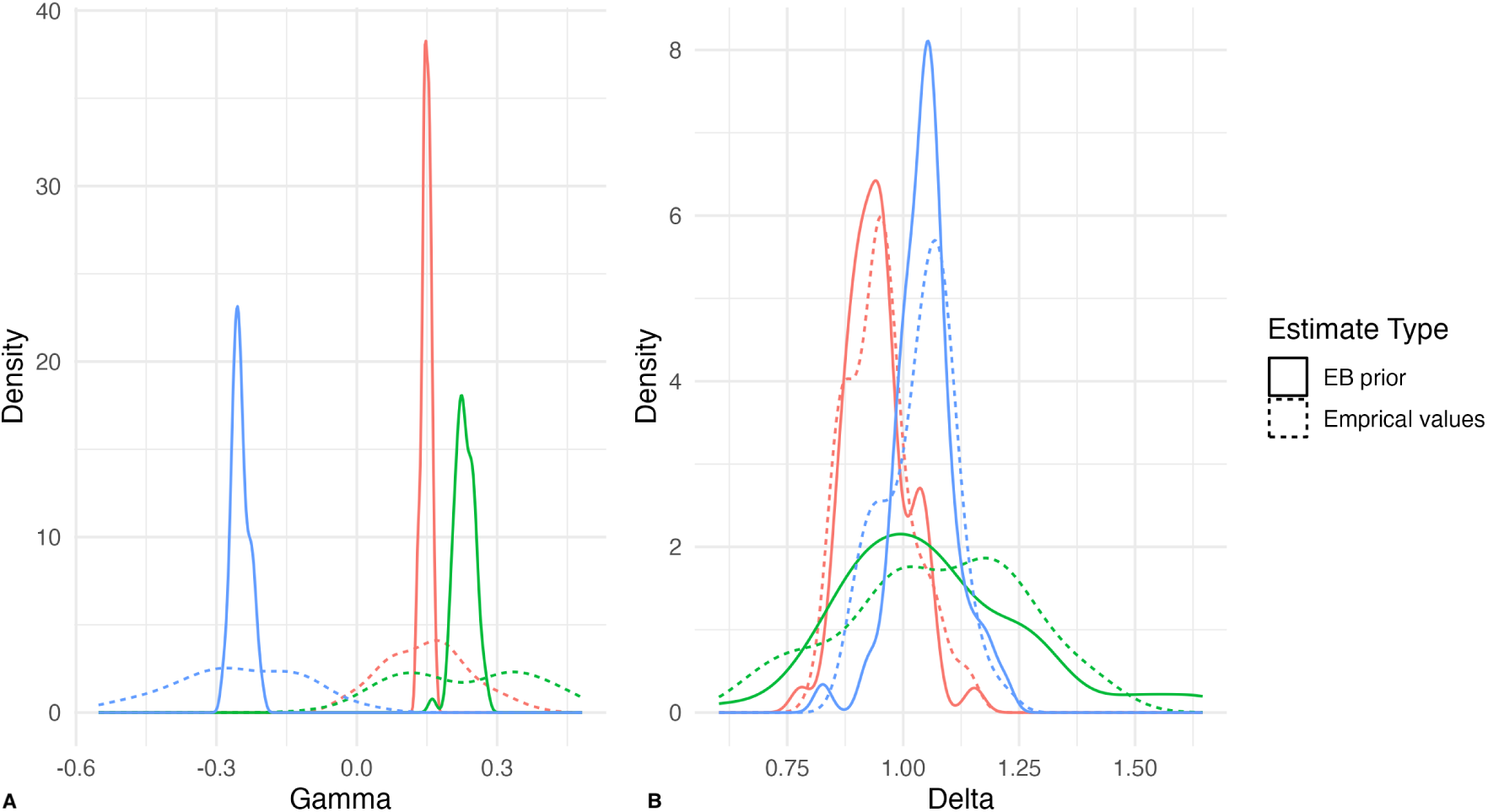
Empirical Bayes diagnostics for longitudinal ComBat. The prior and empirical distributions show close agreement, supporting the use of the empirical Bayes framework.

### 4.3 Harmonization Performance Evaluation

Following Longitudinal ComBat harmonization, the visualizations in Figure 6 indicate substantial improvement in batch alignment. Feature distributions overlap closely across batch levels, and the corresponding age-trend plots show a similarly strong degree of alignment. The residual plots provide no clear evidence of either additive or multiplicative batch effects. Consistent with these findings, both the PCA and *t*-SNE plots show no obvious batch-related clustering, suggesting that global batch effects were substantially reduced.

**Figure 6:**
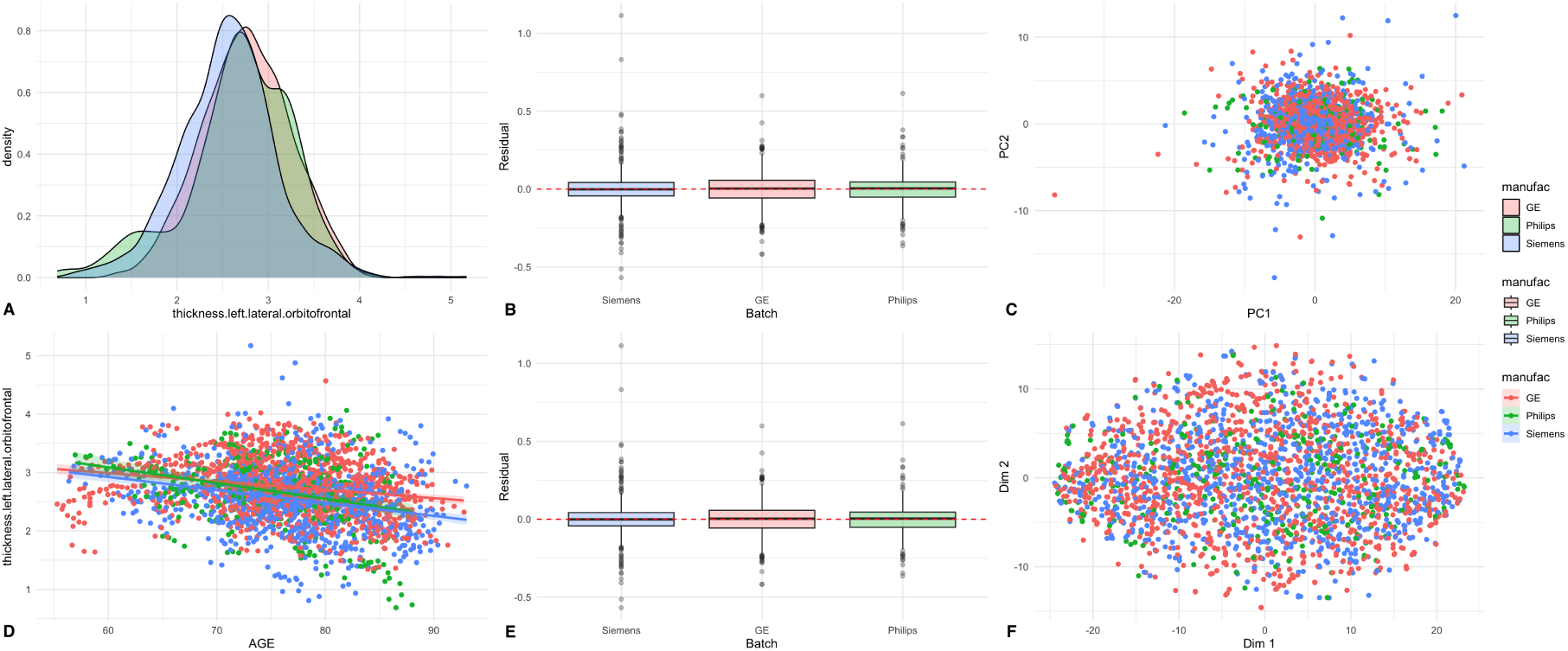
Post-harmonization batch diagnostics. For *thickness.left.lateral.orbitofrontal*, the feature distribution (A) and age-trend plot (D) show substantial overlap across batch levels. The additive residual plot (B) and multiplicative residual plot (E) show minimal remaining batch-related effects for this ROI. The PCA plot (C) and *t*-SNE plot (F) show little visible separation by batch at the multivariate level.

These improvements were also reflected in the statistical test results. Both the Kenward-Roger and ANOVA tests identified 0% of ROIs as significant for additive batch effects, whereas the Kruskal-Wallis test continued to classify 29.03% of ROIs as significant. For multiplicative batch effects, Levene’s test (1.61%) and the Fligner-Killeen test (9.68%) indicated only minor residual heterogeneity across batches, while Bartlett’s test identified no significant results. The MDMR analysis, however, remained significant, which may suggest the presence of subtle global batch effects that are not readily apparent in the PCA or *t*-SNE visualizations.

Overall, these findings suggest that the harmonization procedure effectively removed most feature-level batch effects, although some residual global structure may persist. This result motivates the potential use of CovBat to further address multivariate batch effects.

## 5 Concluding Remarks

As data harmonization has become an important topic and a critical step in data analysis in recent years, particularly in neuroimaging studies, it has become crucial to establish a rigorous harmonization procedure that ensures reproducibility of analytic pipelines and broadens the utility of harmonization models for diverse users and the research community. Despite the development of many harmonization methods, there is currently no systematic pipeline to ensure the smoothness and appropriate use of these methods. A structured and accessible workflow is needed to guide users through diagnosing batch effects, selecting suitable harmonization strategies, and evaluating post-harmonization performance. In practice, researchers often encounter challenges arising from fragmented implementations, inconsistent coding syntax, and limited tools for systematic batch-effect assessment.

Several widely adopted methods, including ComBat (neuroCombat) (Fortin, Cullen, et al. 2018; W. E. Johnson, Li, and Rabinovic 2006), Longitudinal ComBat (Beer et al. 2020), ComBat-GAM (neuroHarmonize) (Pomponio et al. 2020), and CovBat (Chen et al. 2021), provide important statistical frameworks for adjusting mean, variance, or covariance-related batch effects. However, existing implementations primarily focus on model estimation and are typically distributed as standalone tools. Toolbox-based implementations such as DPABI Harmonization (Y.-W. Wang, H.-L. Wang, and Yan 2024) offer graphical interfaces for applying harmonization methods but lack structured and comprehensive diagnostic frameworks for evaluating overall harmonization performance. Applications such as BatchQC (Manimaran et al. 2016) provide Shiny-based visualization and preliminary batch-effect diagnostics, primarily in genomic contexts. However, they are not tailored to neuroimaging data structures and do not provide systematic support for comparing multiple harmonization methods or conducting post-harmonization downstream modeling. In contrast, quality-control platforms such as MRIQC (Esteban, Birman, et al. 2017) and preprocessing pipelines such as fMRIPrep (Esteban, Markiewicz, et al. 2018) emphasize image-level artifact detection rather than statistical harmonization diagnostics. To our knowledge, no neuroimaging-focused software currently integrates batch-effect diagnostics, multiple harmonization strategies, quantitative comparison tools, and downstream modeling support within a unified and reproducible framework.

`ComBatFamQC` was designed to address these gaps by streamlining the harmonization workflow and providing tools for assessing potential batch effects before and after harmonization. Key features of the package include user-friendly batch-effect diagnostics and harmonization through an interactive Shiny app, integration of commonly used harmonization methods using a consistent `R` syntax, and a variety of post-harmonization downstream analyses for users to consider. To accommodate users with diverse coding backgrounds, we also developed two command-line interfaces that allow users to control all stages of the workflow. Detailed tutorials for each stage are available on GitHub at: https://zheng206.github.io/ComBatQC-Web/. By incorporating visualization tools, statistical testing procedures, and downstream analytic modules, `ComBatFamQC` promotes transparent assessment of harmonization performance and supports both in-sample and out-of-sample applications. Its modular architecture also enables straightforward integration of additional harmonization approaches and evaluation tools as the field evolves.

Although `ComBatFamQC` provides useful diagnostics for evaluating potential batch effects, several limitations should be considered when interpreting these results. First, statistical tests for batch-effect detection provide evidence of distributional or multi-variate differences across batches, but they do not, by themselves, quantify the practical importance of those differences. Their sensitivity can depend on sample size, feature dimensionality, group balance, dependence among features, and distributional characteristics. For example, the Kruskal–Wallis test assesses differences in rank distributions across groups and may detect small location or distributional differences in large or unbalanced datasets, even when the magnitude of the batch effect is limited. Similarly, MDMR in `ComBatFamQC` uses Euclidean distance and is therefore sensitive to differences in the overall location, scale, and dispersion of multivariate feature profiles across batches. In large-sample or high-dimensional neuroimaging settings, small residual differences across many regions may accumulate in the Euclidean distance matrix and yield statistically significant results, even when the remaining batch-associated variation is small relative to biological or clinical variation. Therefore, post-harmonization diagnostics should be interpreted alongside visualization, effect-size summaries, covariate balance, and downstream model performance, rather than as standalone indicators of harmonization success or failure.

Second, out-of-sample harmonization relies on assumptions that may not hold equally well in all applications. For the model-based approach, the training data are assumed to provide a representative basis for estimating batch-effect parameters and covariate effects that can be transferred to the target data. This assumption may be weakened when the training sample is small, when important covariates have limited overlap across sites or batches, or when the target data include scanner protocols, demographic distributions, disease characteristics, or acquisition settings that are poorly represented in the training set. Under limited covariate overlap, batch and biological effects may be difficult to distinguish, which can increase the risk of either incomplete batch removal or attenuation of meaningful biological variation. For the reference-based approach, the reference dataset is assumed to be sufficiently homogeneous with respect to batch-related variation, or at least to represent the target scale to which new data should be mapped. If the reference data contain unmodeled batch effects or are not representative of the target population, residual batch effects or unwanted distributional shifts may remain after harmonization. In such settings, out-of-sample harmonization results should be interpreted cautiously, and users are encouraged to examine covariate overlap, batch composition, feature distributions, and post-harmonization diagnostics before drawing substantive conclusions.

Finally, although `ComBatFamQC` integrates several commonly used harmonization methods and diagnostic tools, it does not cover all possible harmonization settings. Future development will focus on expanding the package to accommodate additional harmonization frameworks and more complex study designs, including out-of-sample harmonization for longitudinal ComBat and out-of-sample harmonization within the interactive Shiny workflow. We also plan to allow more flexible covariate selection for lifespan age-trend visualization and broaden the range of post-harmonization downstream analyses. Additional priorities include improving guidance for small-sample settings, developing diagnostics for site-by-covariate confounding, and enhancing assessment of feature dependence and covariance-related batch effects in longitudinal, multimodal, and high-dimensional neuroimaging data.

By streamlining the harmonization process within a unified and accessible platform, `ComBatFamQC` aims to support reproducible, transparent, and adaptable harmonization practices across a wide range of datasets and study designs. In doing so, the package provides both practical tools for applied researchers and an extensible framework for future methodological development, helping to advance harmonization workflows toward greater consistency and collaborative reproducibility within the neuroimaging research community.

## Computational details

The results in this paper were obtained using `R` 4.4.1. `R` and all packages used are available from the Comprehensive `R` Archive Network (CRAN) at https://CRAN.R-project.org/.

## Author Contributions

**Zheng Ren**: Conceptualization, Methodology, Software, Formal Analysis, Investigation, Validation, Data Curation, Visualization, Writing – Original Draft, Writing – Review & Editing.

**Elizabeth A. Horwath**: Software, Validation, Investigation, Visualization, Writing – Review & Editing.

**Siyan Wen**: Software, Validation, Investigation, Visualization, Writing – Review & Editing.

**Randa Melhem**: Validation, Investigation, Writing – Review & Editing.

**Jessica K. Anderson**: Conceptualization, Methodology, Resources, Writing – Review & Editing.

**W. Evan Johnson**: Conceptualization, Methodology, Resources, Writing – Review & Editing.

**Russell T. Shinohara**: Conceptualization, Methodology, Investigation, Validation, Supervision, Project Administration, Funding Acquisition, Writing – Review & Editing.

**Andrew A. Chen**: Conceptualization, Methodology, Software, Resources, Validation, Writing – Review & Editing.

**Haochang Shou**: Conceptualization, Methodology, Investigation, Validation, Supervision, Project Administration, Funding Acquisition, Writing – Review & Editing.

## Acknowledgments

This work was supported by the National Institute of Neurological Disorders and Stroke (U24-NS130411, R01-NS112274), the National Institute of Aging (RF1-AG054409) and the National Institute of Mental Health (R01-MH123550, R01-MH112847). The content is solely the responsibility of the authors and does not necessarily represent the official views of the funding agencies.

Data used in the preparation of this article were obtained from the Alzheimer’s Disease Neuroimaging Initiative (ADNI) database (adni.loni.usc.edu). As such, the investigators within ADNI contributed to the design and implementation of ADNI and/or provided data, but did not participate in the analysis or writing of this report. A complete listing of ADNI investigators can be found at: http://adni.loni.usc.edu/wp-content/uploads/how_to_apply/ADNI_Acknowledgement_List.pdf.

Data collection and sharing for this project was funded by the Alzheimer’s Disease Neuroimaging Initiative (ADNI) (National Institutes of Health Grant U01 AG024904) and DOD ADNI (Department of Defense award number W81XWH-12-2-0012). ADNI is funded by the National Institute on Aging, the National Institute of Biomedical Imaging and Bioengineering, and through generous contributions from the following: AbbVie, Alzheimer’s Association; Alzheimer’s Drug Discovery Foundation; Araclon Biotech; BioClinica, Inc.; Biogen; Bristol-Myers Squibb Company; CereSpir, Inc.; Cogstate; Eisai Inc.; Elan Pharmaceuticals, Inc.; Eli Lilly and Company; EuroImmun; F. Hoffmann-La Roche Ltd and its affiliated company Genentech, Inc.; Fujirebio; GE Healthcare; IXICO Ltd.; Janssen Alzheimer Immunotherapy Research & Development, LLC.; Johnson & Johnson Pharmaceutical Research & Development LLC.; Lumosity; Lundbeck; Merck & Co., Inc.; Meso Scale Diagnostics, LLC.; NeuroRx Research; Neurotrack Technologies; Novartis Pharmaceuticals Corporation; Pfizer Inc.; Piramal Imaging; Servier; Takeda Pharmaceutical Company; and Transition Therapeutics. The Canadian Institutes of Health Research is providing funds to support ADNI clinical sites in Canada. Private sector contributions are facilitated by the Foundation for the National Institutes of Health (www.fnih.org). The grantee organization is the Northern California Institute for Research and Education, and the study is coordinated by the Alzheimer’s Therapeutic Research Institute at the University of Southern California. ADNI data are disseminated by the Laboratory for Neuro Imaging at the University of Southern California.

## Supplementary Material

**Figure S1:**
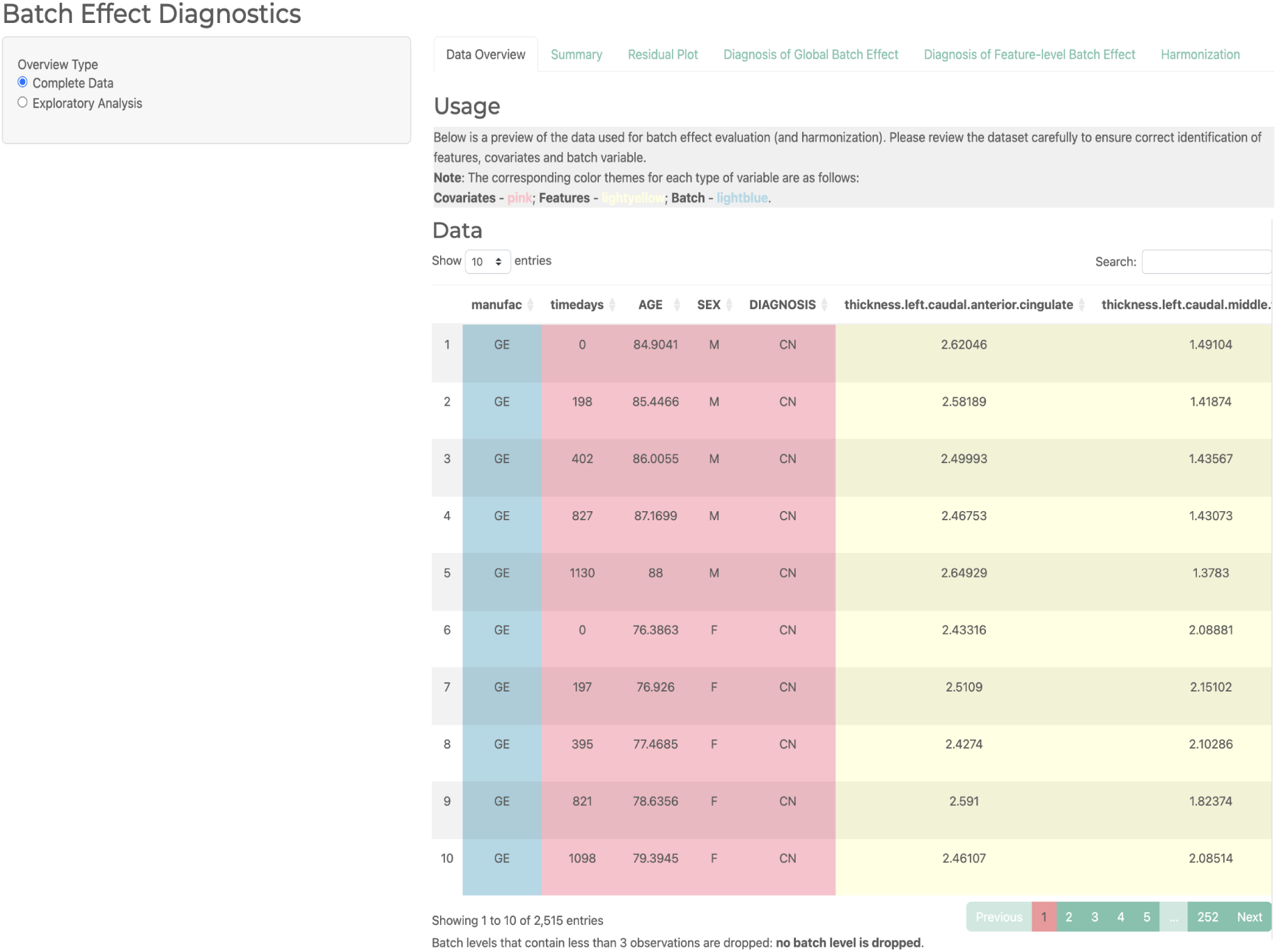
Example of the *Data Overview* section from the interface with *Complete Data* selected. The displayed table represents the dataset used for batch effect diagnostics. Color coding: Blue – batch variable; Pink – covariates; Light yellow – features. A note below the table indicates whether any batch levels were dropped due to insufficient sample size.

**Figure S2:**
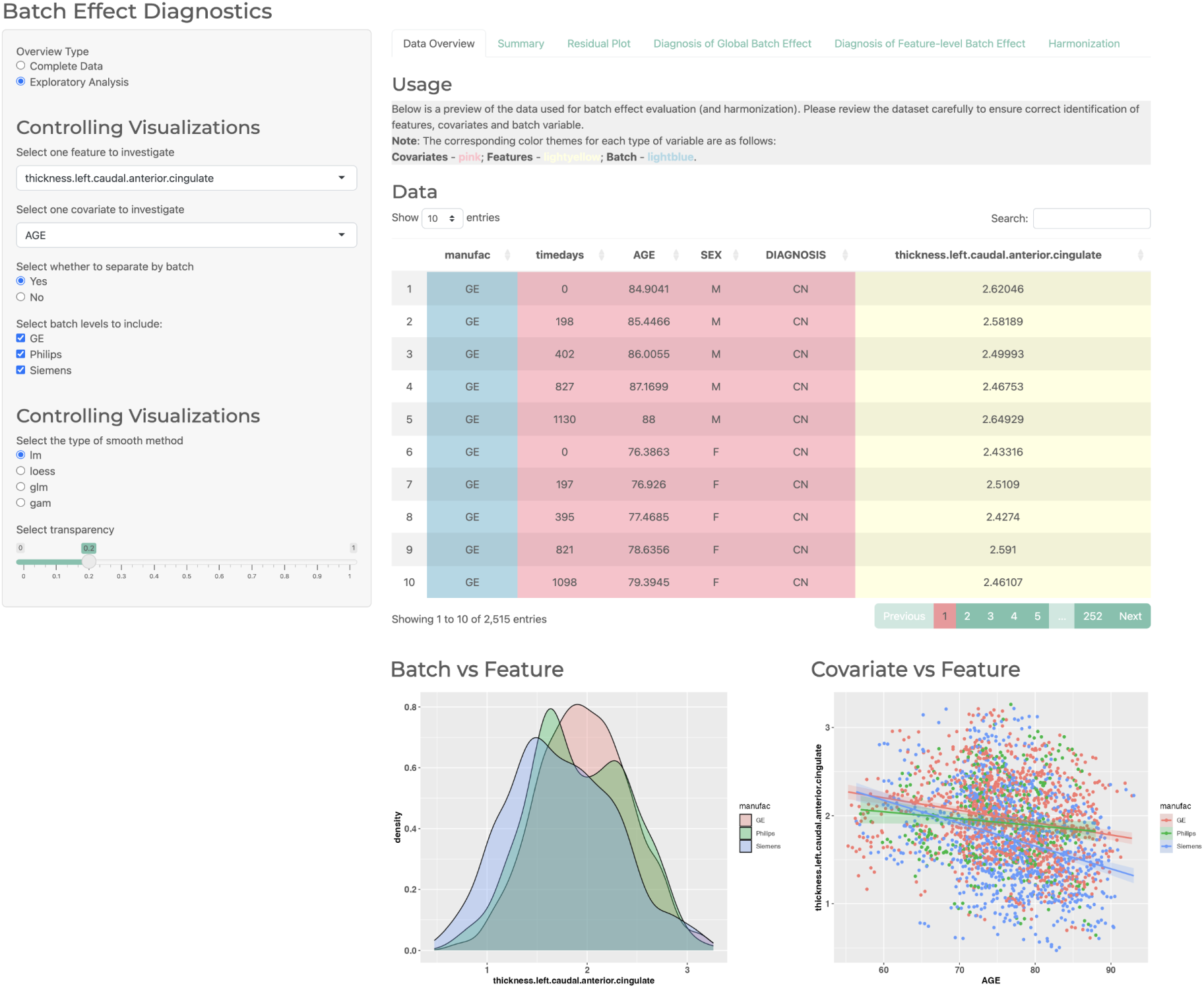
Example of the *Data Overview* section from the interface with *Exploratory Analysis* selected. The Batch vs Feature plot displays density curves of the selected feature across different batch levels, highlighting differences in distributional shape, mean, and scale. Users can choose whether to separate the feature by batch and select which batch levels to include. The Covariate vs Feature plot illustrates the relationship between a selected covariate and feature, allowing users to explore potential non-linear trends. Smoothing methods such as `lm`, `loess`, `glm`, and `gam` can be applied, with adjustable transparency settings.

**Figure S3:**
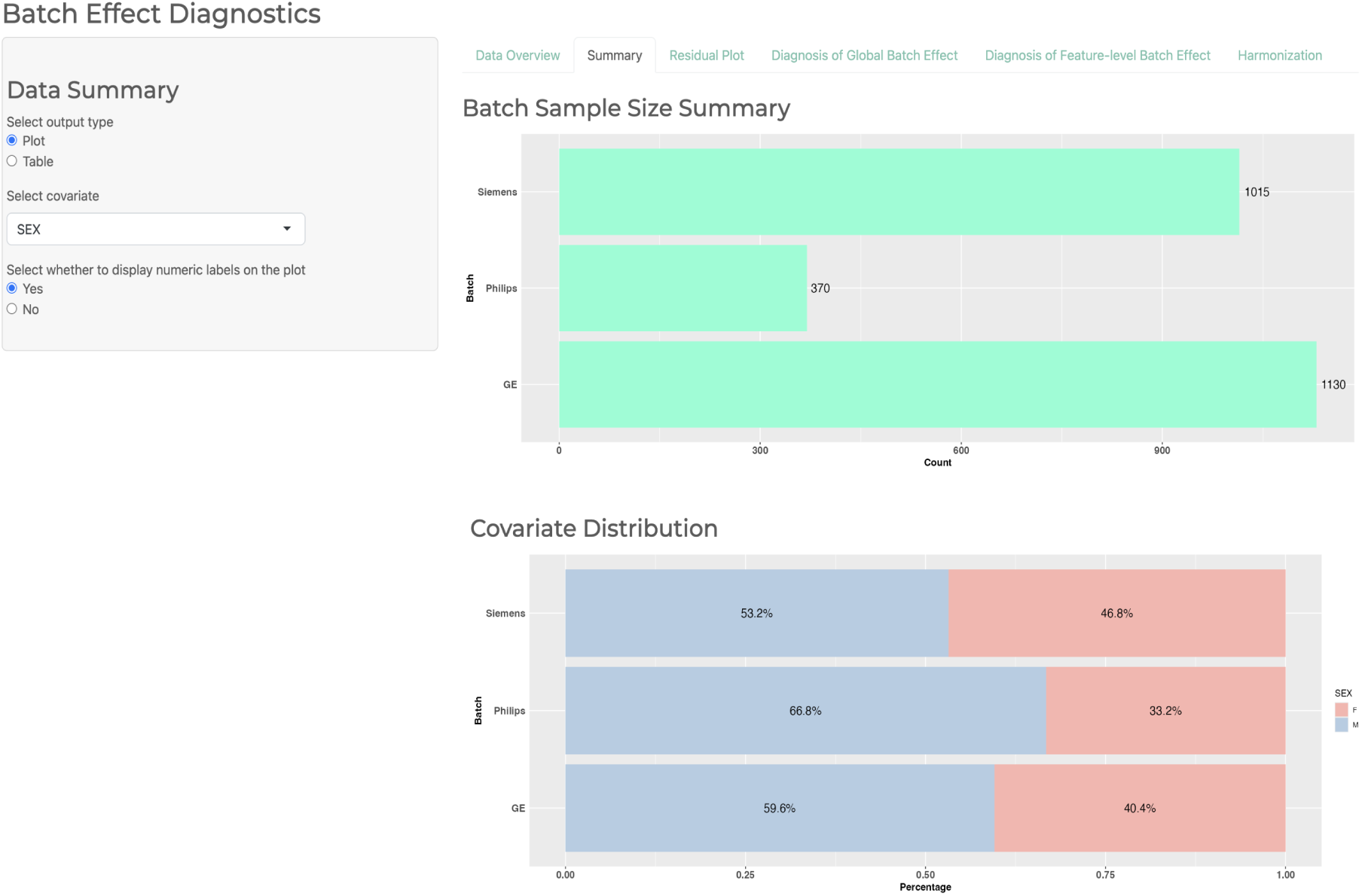
Example of the *Summary* section from the interface with *Plot* selected. The Batch Sample Size Summary plot is a bar chart displaying the sample size for each batch level. The Covariate Distribution plot shows the distribution of the selected covariate across batches. If the covariate is categorical, the plot appears as a stacked bar chart; if numerical, it is displayed as a box plot. Users can choose whether to display numeric labels on the plots.

**Table S1:**
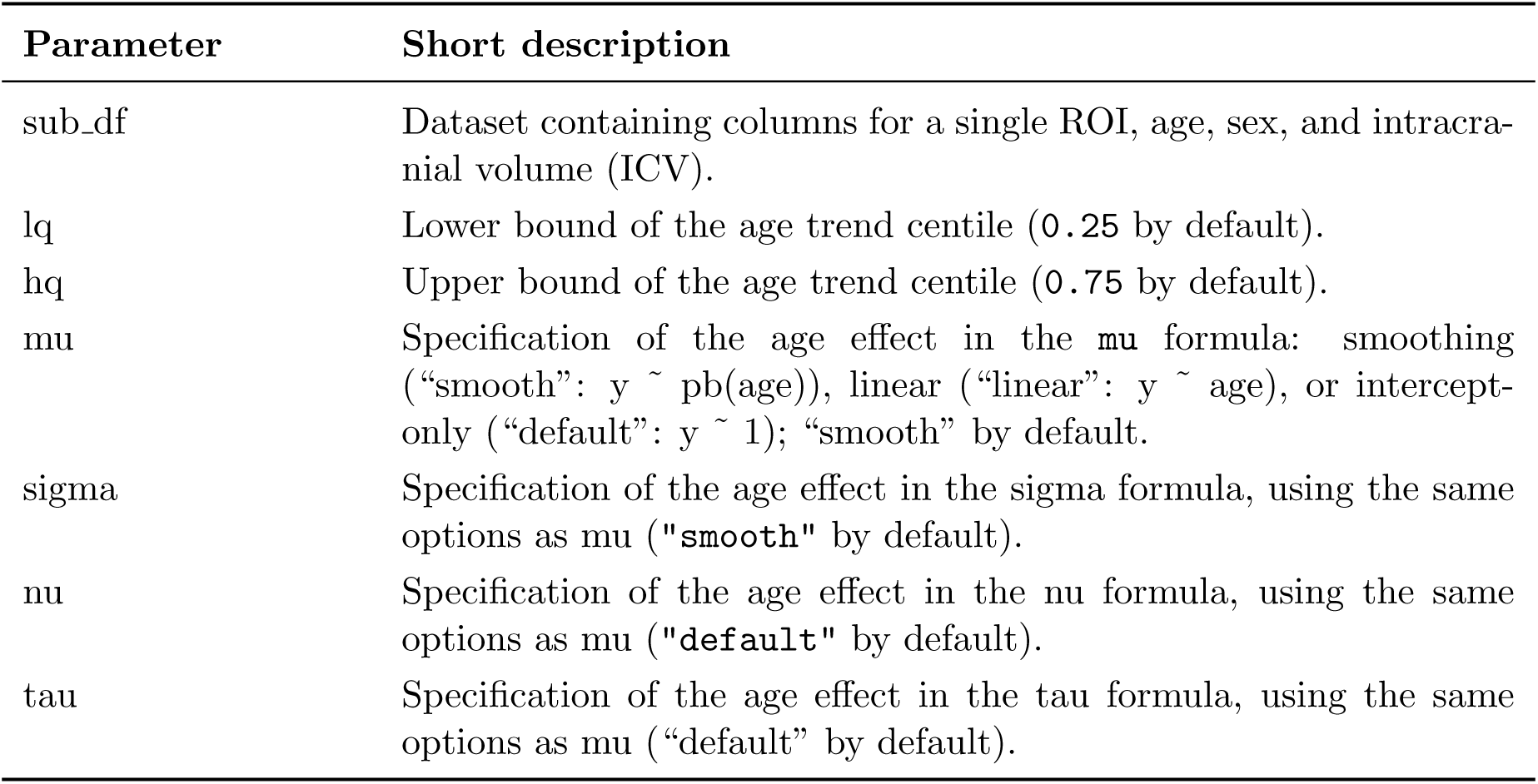
Age trend estimation–related parameters.

| Parameter | Short description |
| --- | --- |
| sub_df | Dataset containing columns for a single ROI, age, sex, and intracranial volume (ICV). |
| lq | Lower bound of the age trend centile (0.25 by default). |
| hq | Upper bound of the age trend centile (0.75 by default). |
| mu | Specification of the age effect in the mu formula: smoothing (“smooth”: $y \sim \text{pb}(\text{age})$), linear (“linear”: $y \sim \text{age}$), or intercept-only (“default”: $y \sim 1$); “smooth” by default. |
| sigma | Specification of the age effect in the sigma formula, using the same options as mu (“smooth” by default). |
| nu | Specification of the age effect in the nu formula, using the same options as mu (“default” by default). |
| tau | Specification of the age effect in the tau formula, using the same options as mu (“default” by default). |

**Figure S4:**
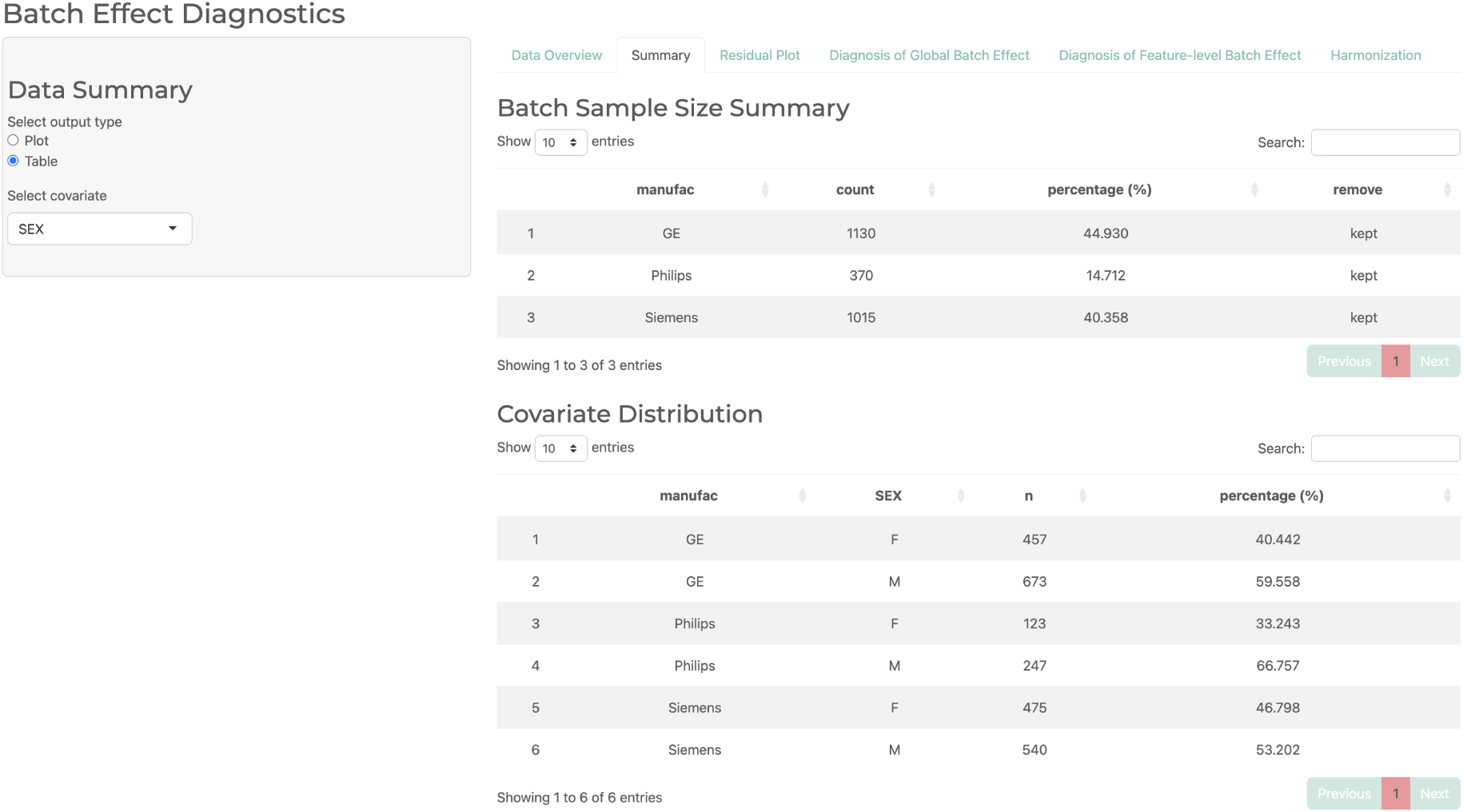
Example of the *Summary* section from the interface with *Table* selected. The Batch Sample Size Summary table displays the count and percentage of samples for each batch level, along with an indicator of whether the batch was removed due to insufficient sample size. The Covariate Distribution table provides summary statistics for the selected covariate, grouped by batch level.

**Figure S5:**
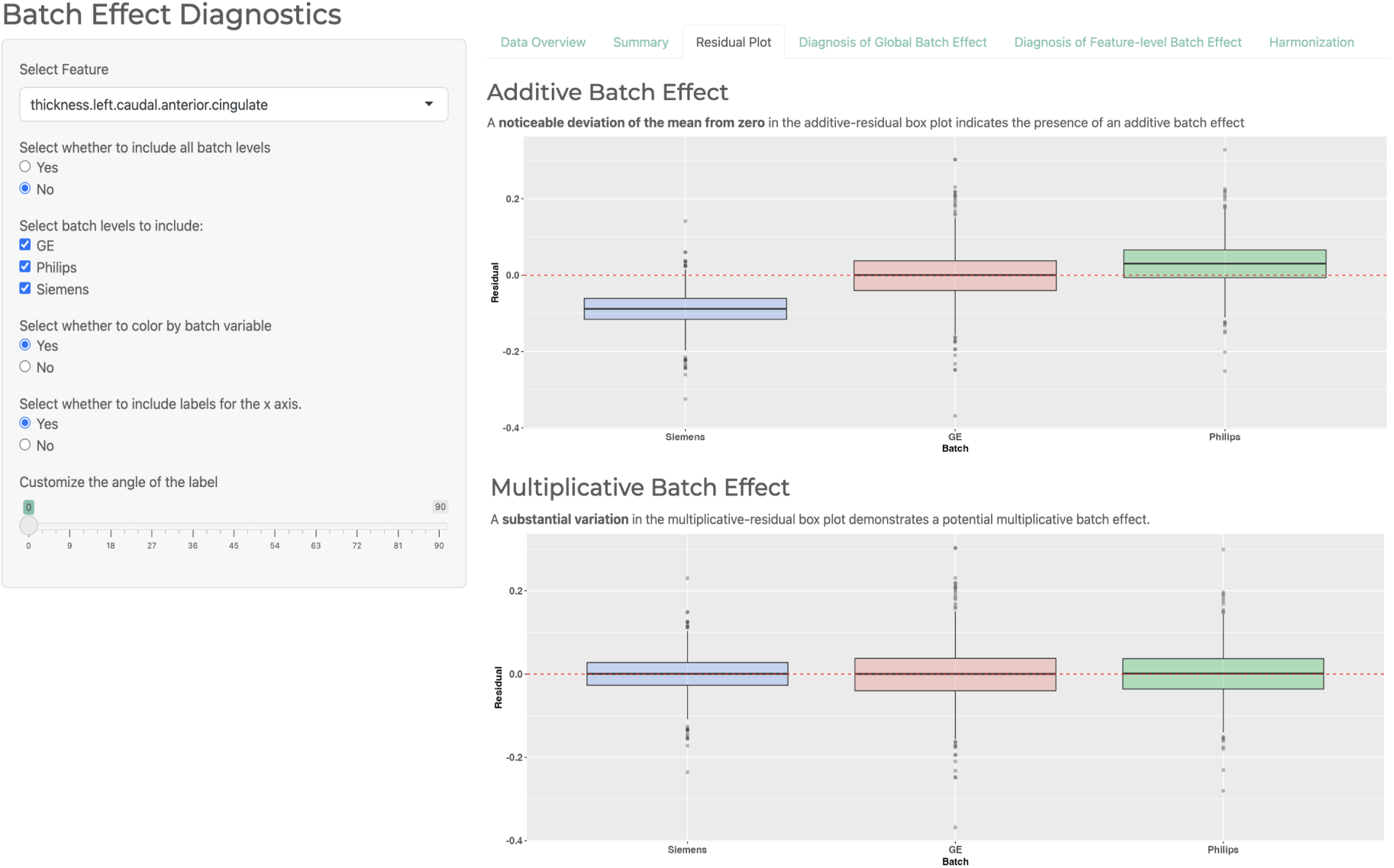
Example of the *Residual Plot* section from the interface. Users can customize the residual plots by selecting specific batch levels, applying color coding by batch, toggling x-axis labels, and adjusting the angle of the x-axis text. The Additive Batch Effect plot highlights deviations in mean residuals across batches, while the Multiplicative Batch Effect plot illustrates differences in residual variance, both indicating potential batch effects.

**Figure S6:**
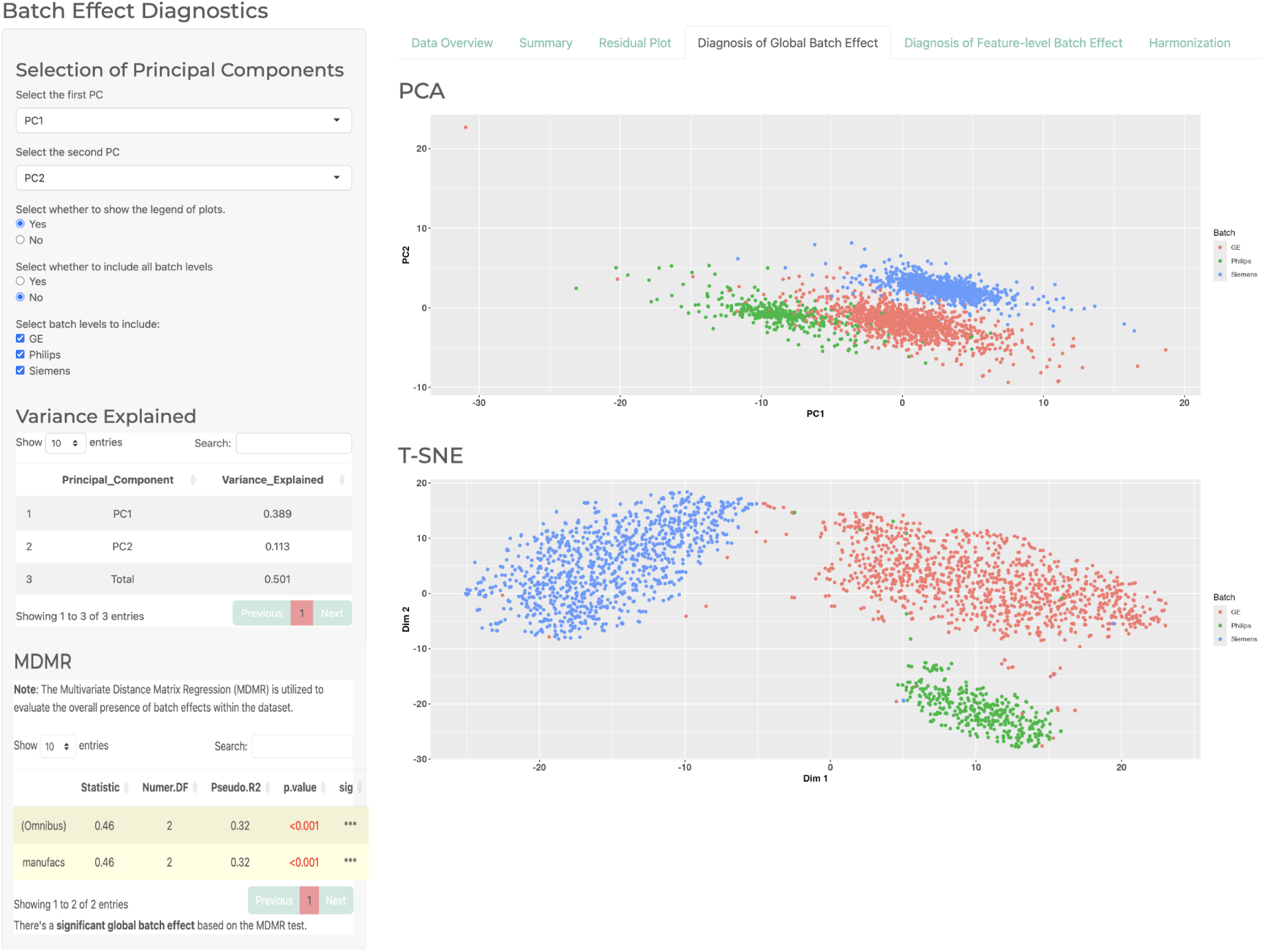
Example of the *Diagnosis of Global Batch Effect* section from the interface. Users can select principal components, toggle the legend display, and choose batch levels to include for better comparison.

**Figure S7:**
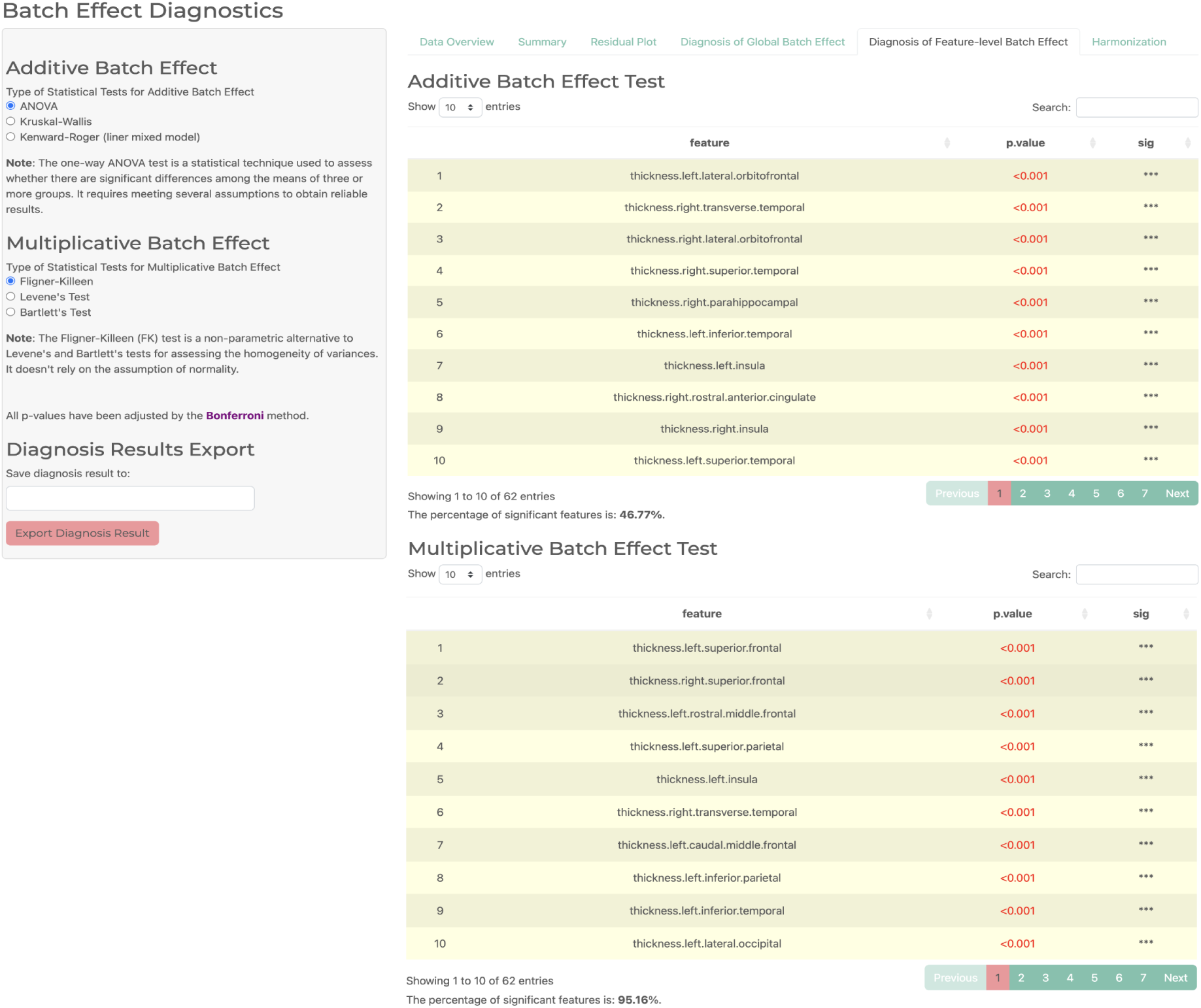
Example of the *Diagnosis of Feature-level Batch Effect* section from the interface. A brief explanation of each method appears once it is selected.

**Figure S8:**
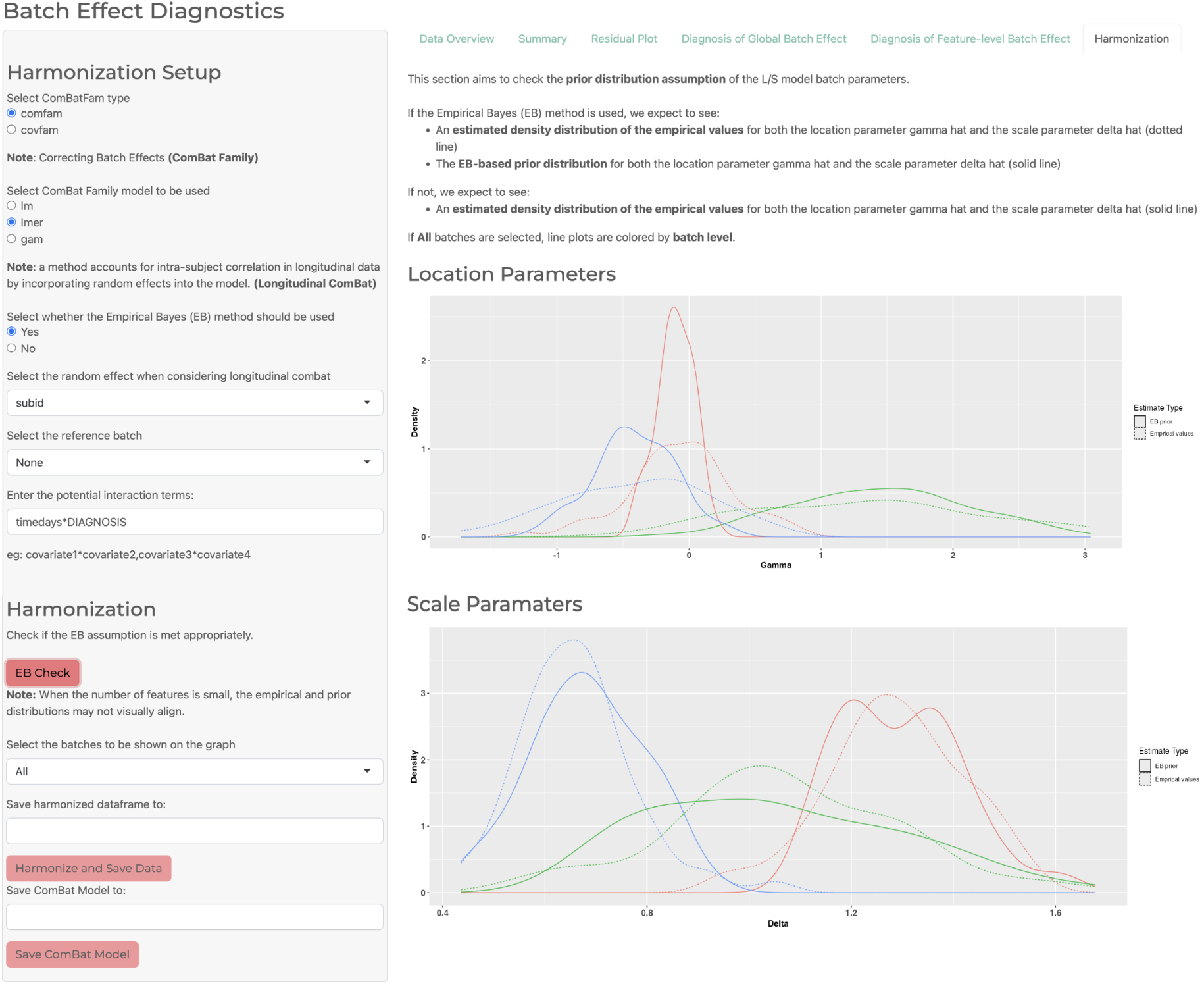
Example of the Interactive *Harmonization* section from the interface. The Location Parameters plot is used to assess whether the normal prior assumption for the location parameter (gamma) holds, while the Scale Parameters plot evaluates the inverse gamma prior assumption for the scale parameter (delta). When Empirical Bayes (EB) is used, both the empirical (dotted line) and EB-based prior (solid line) distributions are shown for each batch. Users can also select individual batch levels for further investigation.

**Figure S9:**
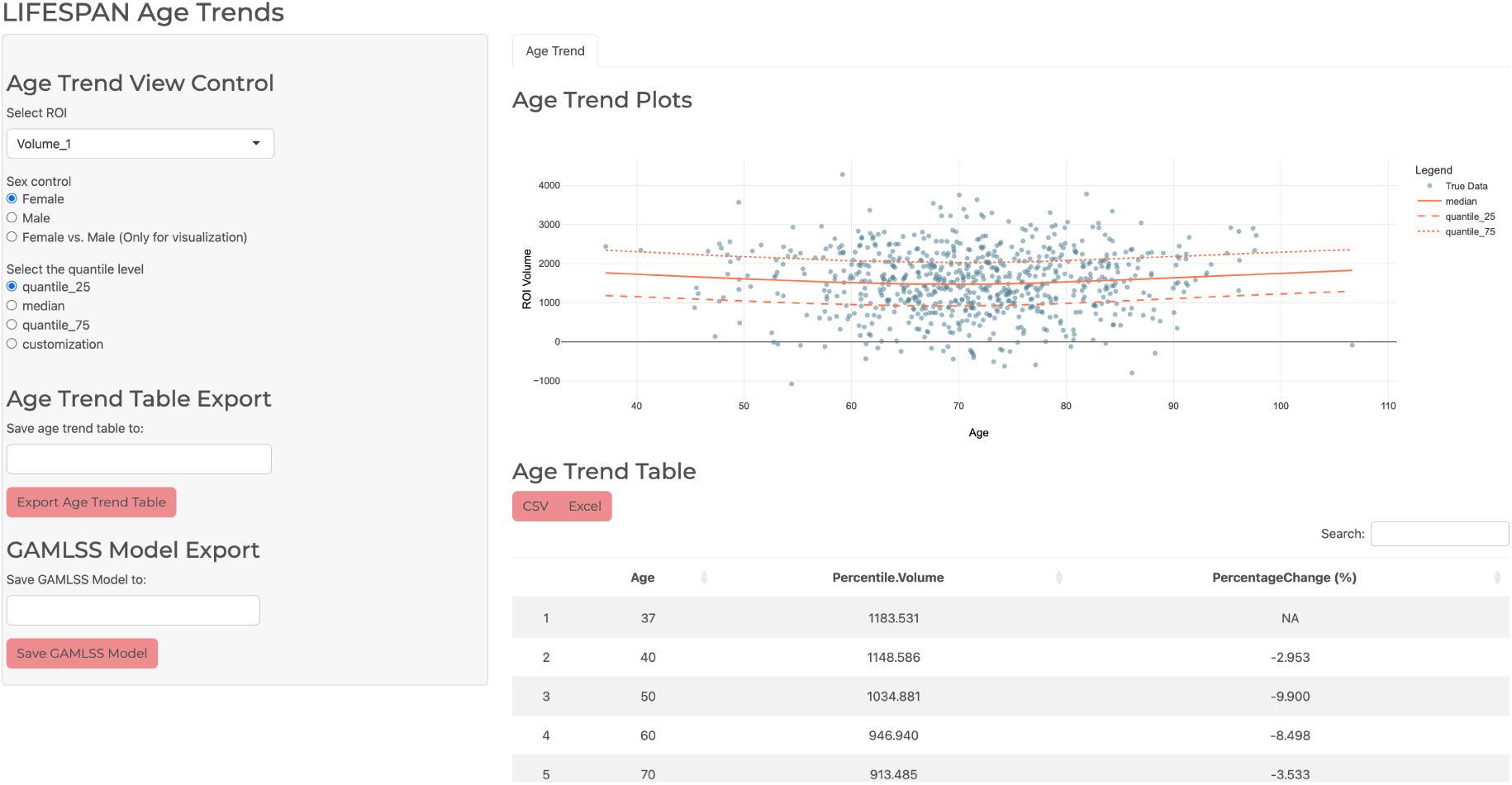
Brain feature lifespan age trend visualization through the Shiny app.

## References

Bartlett, Maurice Stevenson (1937). “Properties of sufficiency and statistical tests”. In: Proceedings of the Royal Society of London. Series A-Mathematical and Physical Sciences 160.901, pp. 268–282.

Bates, Douglas et al. (2015). “Fitting Linear Mixed-Effects Models Using lme4”. In: Journal of Statistical Software 67.1, pp. 1–48. doi: 10.18637/jss.v067.i01.

Bayer, Johanna M. M., et al. (Oct. 2022). “Site effects how-to and when: An overview of retrospective techniques to accommodate site effects in multi-site neuroimaging analyses”. In: Frontiers in Neurology 13. issn: 1664-2295. doi: 10.3389/fneur.2022.923988. url: 10.3389/fneur.2022.923988.

Beer, Joanne C., et al. (Oct. 2020). “Longitudinal ComBat: A method for harmonizing longitudinal multi-scanner imaging data”. In: NeuroImage 220, p. 117129. doi: 10.1016/j.neuroimage.2020.117129. url: 10.1016/j.neuroimage.2020.117129.

Chen, Andrew A., et al. (Dec. 2021). “Mitigating site effects in covariance for machine learning in neuroimaging data”. In: Human Brain Mapping 43.4, pp. 1179–1195. issn: 1097-0193. doi: 10.1002/hbm.25688. url: 10.1002/hbm.25688.

Conover, William J, Mark E Johnson, and Myrle M Johnson (1981). “A comparative study of tests for homogeneity of variances, with applications to the outer continental shelf bidding data”. In: Technometrics 23.4, pp. 351–361.

Esteban, Oscar, Daniel Birman, et al. (Sept. 2017). “MRIQC: Advancing the automatic prediction of image quality in MRI from unseen sites”. In: PLOS ONE 12.9. Ed. by Boris C Bernhardt, e0184661. issn: 1932-6203. doi: 10.1371/journal.pone.0184661. url: 10.1371/journal.pone.0184661.

Esteban, Oscar, Christopher J. Markiewicz, et al. (Dec. 2018). “fMRIPrep: a robust preprocessing pipeline for functional MRI”. In: Nature Methods 16.1, pp. 111–116. issn: 1548-7105. doi: 10.1038/s41592-018-0235-4. url: 10.1038/s41592-018-0235-4.

Fortin, Jean-Philippe, Nicholas Cullen, et al. (Feb. 2018). “Harmonization of cortical thickness measurements across scanners and sites”. In: NeuroImage 167, pp. 104–120. issn: 1053-8119. doi: 10.1016/j.neuroimage.2017.11.024. url: 10.1016/j.neuroimage.2017.11.024.

Fortin, Jean-Philippe, Drew Parker, et al. (Nov. 2017). “Harmonization of multi-site diffusion tensor imaging data”. In: NeuroImage 161, pp. 149–170. issn: 1053-8119. doi: 10.1016/j.neuroimage.2017.08.047. url: 10.1016/j.neuroimage.2017.08.047.

Fox, John and Sanford Weisberg (2019). An R Companion to Applied Regression. Third. Thousand Oaks CA: Sage. url: https://socialsciences.mcmaster.ca/jfox/Books/Companion/.

Halekoh, Ulrich and Søren Højsgaard (2014). “A Kenward-Roger Approximation and Parametric Bootstrap Methods for Tests in Linear Mixed Models – The R Package pbkrtest”. In: Journal of Statistical Software 59.9, pp. 1–30. url: https://www.jstatsoft.org/v59/i09/.

Hotelling, H. (Sept. 1933). “Analysis of a complex of statistical variables into principal components.” In: Journal of Educational Psychology 24.6, pp. 417–441. doi: 10.1037/h0071325. url: 10.1037/h0071325.

Hu, Fengling et al. (July 2023). “Image harmonization: A review of statistical and deep learning methods for removing batch effects and evaluation metrics for effective harmonization”. In: NeuroImage 274, p. 120125. issn: 1053-8119. doi: 10.1016/j.neuroimage.2023.120125. url: 10.1016/j.neuroimage.2023.120125.

Jack, Clifford R., et al. (Feb. 2008). “The Alzheimer’s disease neuroimaging initiative (ADNI): MRI methods”. In: Journal of Magnetic Resonance Imaging 27.4, pp. 685–691. doi: 10.1002/jmri.21049. url: 10.1002/jmri.21049.

Johnson, W. Evan, Cheng Li, and Ariel Rabinovic (Apr. 2006). “Adjusting batch effects in microarray expression data using empirical Bayes methods”. In: Biostatistics 8.1, pp. 118–127. issn: 1465-4644. doi: 10.1093/biostatistics/kxj037. url: 10.1093/biostatistics/kxj037.

Kenward, Michael G and James H Roger (1997). “Small sample inference for fixed effects from restricted maximum likelihood”. In: Biometrics, pp. 983–997.

Kruskal, William H. and W. Allen Wallis (Dec. 1952). “Use of Ranks in One-Criterion Variance Analysis”. In: Journal of the American Statistical Association 47.260, pp. 583–621. issn: 1537-274X. doi: 10.1080/01621459.1952.10483441. url: 10.1080/01621459.1952.10483441.

Levene, Howard et al. (1960). “Contributions to probability and statistics”. In: Essays in honor of Harold Hotelling 278, p. 292.

Manimaran, Solaiappan, et al. (Aug. 2016). “BatchQC: interactive software for evaluating sample and batch effects in genomic data”. In: Bioinformatics 32.24, pp. 3836–3838. issn: 1367-4811. doi: 10.1093/bioinformatics/btw538. url: 10.1093/bioinformatics/btw538.

McArdle, Brian H. and Marti J. Anderson (Jan. 2001). “FITTING MULTIVARIATE MODELS TO COMMUNITY DATA: A COMMENT ON DISTANCE-BASED REDUNDANCY ANALYSIS”. In: Ecology 82.1, pp. 290–297. issn: 0012-9658. doi: 10.1890/0012-9658(2001)082[0290:fmmtcd]2.0.co;2. url: 10.1890/0012-9658(2001)082[0290:FMMTCD]2.0.CO;2.

McArtor, Daniel B., Gitta H. Lubke, and C. S. Bergeman (Oct. 2016). “Extending multivariate distance matrix regression with an effect size measure and the asymptotic null distribution of the test statistic”. In: Psychometrika 82.4, pp. 1052–1077. doi: 10.1007/s11336-016-9527-8. url: 10.1007/s11336-016-9527-8.

Pomponio, Raymond et al. (Mar. 2020). “Harmonization of large MRI datasets for the analysis of brain imaging patterns throughout the lifespan”. In: NeuroImage 208, p. 116450. issn: 1053-8119. doi: 10.1016/j.neuroimage.2019.116450. url: 10.1016/j.neuroimage.2019.116450.

R Core Team (2022). R: A Language and Environment for Statistical Computing. R Foundation for Statistical Computing. Vienna, Austria. url: https://www.R-project.org/.

Rigby, R. A. and D. M. Stasinopoulos (Apr. 2005). “Generalized Additive Models for Location, Scale and Shape”. In: Journal of the Royal Statistical Society Series C: Applied Statistics 54.3, pp. 507–554. issn: 1467-9876. doi: 10.1111/j.1467-9876.2005.00510.x. url: 10.1111/j.1467-9876.2005.00510.x.

van der Maaten, L.J.P. and G.E. Hinton (2008). “Visualizing High-Dimensional Data Using t-SNE”. In: Journal of Machine Learning Research 9, pp. 2579–2605.

Wang, Yu-Wei, Han-Lin Wang, and Chao-Gan Yan (2024). “DPABI harmonization: A toolbox for harmonizing multi-site brain imaging for big-data era”. In: Imaging Neuroscience 2. issn: 2837-6056. doi: 10.1162/imag_a_00388. url: 10.1162/imag_a_00388.

Wickham, Hadley (2016). ggplot2: Elegant Graphics for Data Analysis. Springer-Verlag New York. isbn: 978-3-319-24277-4. url: https://ggplot2.tidyverse.org.

Wood, S. N. (2011). “Fast stable restricted maximum likelihood and marginal likelihood estimation of semiparametric generalized linear models”. In: Journal of the Royal Statistical Society (B) 73.1, pp. 3–36.

Xin, Yao et al. (May 2026). “ComBat-Predict Enhances Generalizability of Neuroimaging Models to New Sites”. In: Human Brain Mapping 47.8. issn: 1097-0193. doi: 10.1002/hbm.70546. url: 10.1002/hbm.70546.

